# The majority of the matrix protein TapA is dispensable for biofilm formation by *Bacillus subtilis*

**DOI:** 10.1101/794164

**Authors:** Chris Earl, Sofia Arnaouteli, Tetyana Sukhodub, Cait E. MacPhee, Nicola R. Stanley-Wall

## Abstract

Biofilm formation is a co-operative behaviour where microbial cells become embedded in an extracellular matrix. This biomolecular matrix plays a key role in the manifestation of the beneficial or detrimental outcome mediated by the collective of cells. *Bacillus subtilis* is an important bacterium for understanding the general principles of biofilm formation and is a plant growth-promoting organism. The protein components of the *B. subtilis* matrix include the secreted proteins BslA, which forms a hydrophobic coat over the biofilm, and TasA, which forms protease-resistant fibres needed for structuring. A third protein TapA (for <u>T</u>asA <u>a</u>nchoring and assembly <u>p</u>rotein) is needed for biofilm formation and helps TasA fibre formation *in vivo* but is dispensable for TasA-fibre assembly *in vitro*. Here we show that TapA is subjected to proteolytic cleavage in the biofilm and that only the first 57 amino acids of the 253-amino acid protein are required for biofilm architecture. However, through the construction of a strain which lacks all eight extracellular proteases, we show that proteolytic cleavage by these enzymes is not a prerequisite for TapA function. It remains unknown why TapA is synthesized at a full length of 253 amino acids when the first 57 are sufficient for biofilm structuring, but the findings do not exclude the core conserved region of TapA having a second role beyond that of structuring the *B. subtilis* biofilm.

## Introduction

The predominant state in which bacteria and archaea live on Earth is in the form of biofilms (Flemming *et al.*, 2016): aggregates of microorganisms embedded in a self-made extracellular matrix. Molecules that can be found in the biofilm matrix include extracellular DNA, exopolysaccharides, lipids and proteins, the latter of which can self-assemble into a variety of functional forms including filaments, films, or fibres (Erskine *et al.*, 2018a). The biofilm matrix conveys ‘emergent properties’ to the cells in the biofilm (Dragos & Kovacs, 2017) including, but not limited to, providing structure and stability to the community (Hobley *et al.*, 2015), aiding the sequestration of nutrients and retaining extracellular enzymes which facilitates further processing of the biofilm matrix (Flemming et al., 2016).

Biofilm formation by the Gram-positive bacterium *Bacillus subtilis* has been intensively studied and the production of the molecules in the matrix is known to be highly controlled by a suite of transcription regulators (Cairns *et al.*, 2014). The exopolymeric matrix of the *B. subtilis* biofilm is comprised of polymers that include an exopolysaccharide (EPS) and an amphiphilic protein BslA which assembles to form a hydrophobic film on the biofilm surface (Kobayashi & Iwano, 2012, Hobley *et al.*, 2013). The major protein component of the *Bacillus subtilis* matrix is protease-resistant fibres formed by the secreted protein TasA (Erskine *et al.*, 2018b, Romero *et al.*, 2010). These fibres provide structural integrity to the biofilm and are necessary for the characteristic wrinkled phenotype of biofilms and pellicles (Branda *et al.*, 2006).

The *tasA* coding region is located within an operon alongside two other genes: namely *sipW* and *tapA* (formerly *yqxM*) (Zhu & Stulke, 2018). SipW is a specialised signal peptidase that is linked with removal of the signal peptide from both TasA (Stover & Driks, 1999b) and TapA (Stover & Driks, 1999a) during the secretion process. SipW also has links with the control of expression of the *epsA-O* operon which encodes the proteins needed for the biofilm exopolysaccharide (Terra *et al.*, 2012). Thus, *sipW* is essential for biofilm formation. TapA is described as an accessory protein that is needed for the formation of TasA fibres (Romero *et al.*, 2011, Romero et al., 2010), and more specifically, for the attachment of the TasA fibres to the cell surface. Secondary structure analysis revealed TapA to be a two-domain protein with significant regions of disorder (Abbasi *et al.*, 2019). Additionally TapA has recently been shown to form fibres (El Mammeri *et al.*, 2019). The absence of TapA is correlated with a reduction in the level of TasA in the biofilm matrix (Romero *et al.*, 2014). Evidence indicates however that, in the absence of TapA, recombinant TasA self-assembles into protease-resistant fibres that are structurally and functionally comparable to native fibres extracted from *B. subtilis* (El Mammeri et al., 2019, Erskine et al., 2018b). Moreover, when provided exogenously these self-assembled recombinant TasA fibres are also biologically active *in vivo* in the absence of TapA (Erskine et al., 2018b). Thus, further evaluation of the function and activity of TapA is warranted.

Here we identify that amino acids 1-57 (inclusive) of TapA form a minimal functional unit of the protein that is required to give rise to the complex architecture of the *B. subtilis* biofilm. Ectopic provision of the DNA encoding this truncated form of TapA is sufficient to restore rugose biofilm formation to the *tapA* deletion strain (the full length protein is 253 amino acids in length). We identify, through site directed mutagenesis, key amino acids in the minimal, functional TapA form that are required for bioactivity and, in doing so, uncover essential hydrophobic amino acids. We show that *in vivo* TapA is proteolytically cleaved to lower molecular weight forms by the native extracellular proteases secreted into the external environment. Finally we establish that proteolysis of TapA by the extracellular proteases is not a prerequisite to activity, suggesting that TapA is degraded after it has fulfilled its function *in vivo*.

## Results

### Construction and complementation of a *B. subtilis tapA* mutant

To investigate the functional regions of *tapA*, an in-frame *tapA* deletion strain was constructed and the deleterious impact on biofilm formation confirmed (Romero et al., 2011) (Fig. 1A). As expected, the rugosity exhibited by the biofilm formed by the parental NCIB3610 isolate was absent when *tapA* was deleted, and was fully reinstated when the *tapA* coding region was expressed from the ecotopic *amyE* locus under the control of the IPTG inducible promoter, Pspank (Fig. 1A). For comparison purposes, a *tasA* deletion strain was examined. This strain also formed flat biofilms that lacked the corrugations and ridges displayed by the wild type strain (Fig. 1A). However, subtle, but consistent differences between the morphology of the *tapA* and *tasA* deletion strains were apparent, with the *tapA* deletion strain occupying a larger footprint than the *tasA* deletion strain (Fig. 1A).

**Figure 1:**
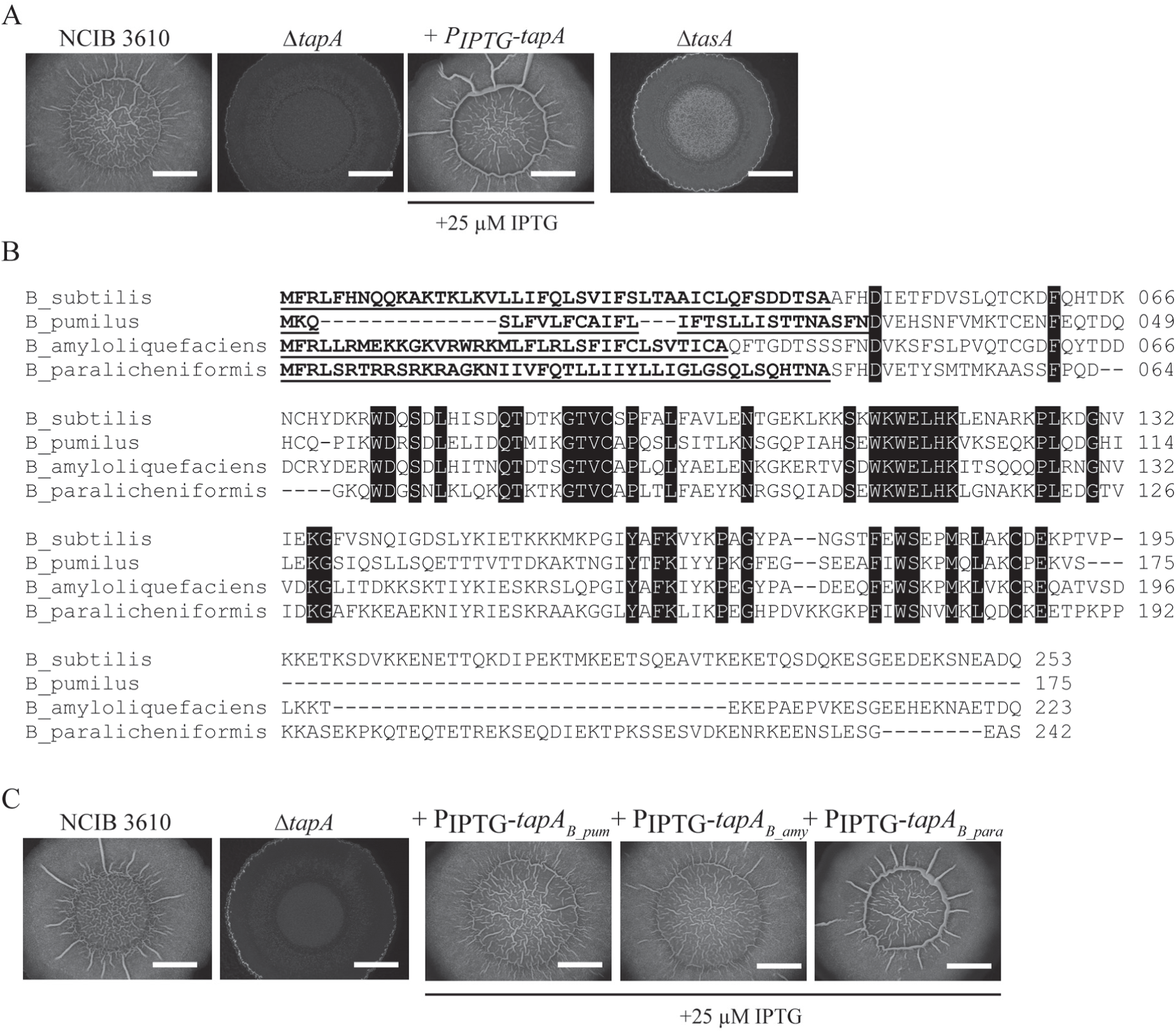
The *tapA* mutant can be complemented by orthologous genes. (**A**) Biofilms formed by NCIB3610, Δ*tapA* (NRS3936), +P_IPTG_-*tapA* (NRS5045) and Δ*tasA* (NRS5267). n> 3; (**B**) Alignment of TapA protein sequences from *B. subtilis, B. pumilus, B. amyloliquefaciens* and *B. paralicheniformis*. The percentage amino acid sequence identity with regards the *B. subtilis* TapA sequence is as follows: *B. pumilus*- 42%, *B. amyloliquefaciens*- 49% and *B. paralicheniformis*- 38%. The bold underlined sequence represents the signal sequence, the black boxes indicate identical amino acids; (**C**) Biofilms formed by NCIB3610, Δ*tapA* (NRS3936), +P_IPTG_-*tapA*_*B_pum*_ (NRS5046), +P_IPTG_-*tapA*_*B_amy*_ (NRS5047), and +P_IPTG_-*tapA*_*B_para*_ (NRS5741). n > 3. Biofilms were grown at 30°C for 48 hours in the presence of IPTG as indicated. The scale bars represent 1 cm.

The *tapA* gene is present in a range of *Bacillus* species and analysis of the protein sequences of TapA from *B. subtilis, B. amyloliquefaciens, B. paralicheniformis* and *B. pumilus* reveals domains with a high degree of conservation and other regions of variability. There are also marked differences in the length of the *tapA* coding region between the different species (Fig. 1B) (Romero et al., 2014). We tested whether these variations in gene length impacted the ability of the orthologous coding regions to genetically substitute for the *B. subtilis tapA* coding region. Heterologous expression of the *tapA* coding region from *B. amyliquofacienes, B. paralicheniformis* and *B. pumilus* in the *B. subtilis tapA* deletion strain resulted in IPTG-dependent recovery of biofilm architecture, such that the resulting biofilms were indistinguishable from those formed by the wild type NCIB3610 *B. subtilis* strain (Fig. 1C). These findings are consistent with, and extend, those previously published that revealed *tapA* from *B. amyliquofacienes* was able to functionally replace TapA of *B. subtilis* (Romero et al., 2014), indicating the presence of conserved amino acids necessary for TapA function.

### Identification of a TapA minimal functional unit

Previous work has preliminarily explored if TapA has a functional unit that is shorter than the full length coding region: the coding region for amino acids 194-230 of TapA could be deleted and the resulting truncated protein retained activity (Romero et al., 2014). Therefore, to determine the length of the *tapA* minimal functional coding region, we systematically constructed truncations of the *tapA* coding region, deleting the *tapA* coding region sequentially from the 3’ end. The ability of the variant-length *tapA* constructs to genetically complement the *tapA* deletion strain was tested after integration to the chromosome at the *amyE* locus. Expression of the coding region was placed under the control of the IPTG inducible promoter. In total 22 variants of the *tapA* coding region were assessed (Fig. 2A, S1A). We noted that the TapA_1-60_ variant was fully functional while the TapA_1-50_ form lacked activity and could not restore biofilm architecture to the *tapA* mutant (Fig 2B). Therefore we made additional constructs with finer scale changes in the region coding amino acids 60 and 50 of TapA (Fig 2C). Through this analysis it was established that amino acids 1-57, inclusive, represent the minimal form of TapA that is capable of reinstating complex biofilm formation to the *tapA* deletion strain. This conclusion was reached primarily through a visual analysis of biofilm rugosity (Fig. 2B and 2C, S1A). Additionally, as the level of TasA (25.7 kDa) in the biofilm is substantially reduced in the absence of functional TapA (Romero et al., 2014) (Fig. S1B), the bioactivity of the truncations were also supported by immunoblot analysis which showed a recovery of TasA levels (Fig. S1B). The identification of this region of *tapA* as sufficient for TapA activity is consistent with the previous identification of amino acids 50-57 as being needed for TapA function in the full length protein (Romero et al., 2014).

**Figure 2:**
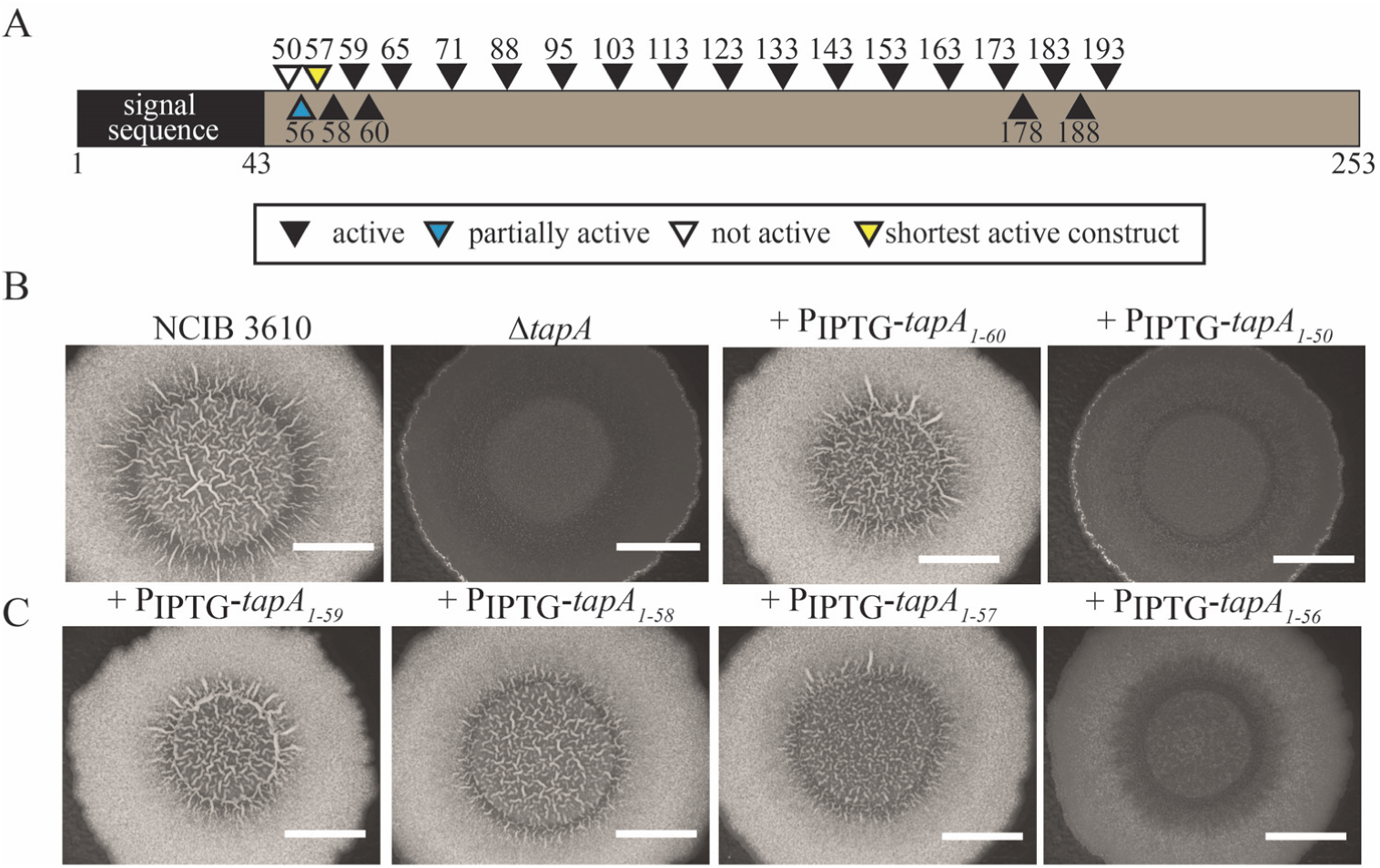
The *tapA* coding region is functional when truncated. (**A**) Schematic of the constructs generated and ability to restore rugose biofilm formation to the Δ*tapA* deletion strain when expressed from an ectopic position on the chromosome. Drawn approximately to scale.; (**B**) Biofilms formed by NCIB3610, Δ*tapA* (NRS3936), +P_IPTG_-*tapA*_*1-60*_ (NRS6044), and *tapA*_*1-50*_ (NRS6002); (**C**) Biofilms formed by +P_IPTG_-*tapA*_*1-59*_ (NRS6043), +P_IPTG_-*tapA*_*1-58*_ (NRS6042), +P_IPTG_-*tapA*_*1-57*_ (NRS6041), and +P_IPTG_-*tapA*_*1-56*_ (NRS6025). Biofilms were grown at 30°C for 48 hours in the presence of 25 µM IPTG. Biofilm images are representative of at least 3 independent replicates. The scale bars represent 1 cm.

### Amino acids critical for function in the minimal functional region of TapA

We were interested in the features of the TapA minimal form that conferred activity. The predicted TapA signal sequence likely comprises 43 amino acids (Petersen *et al.*, 2011) (not 33 amino acids as previously stated (Romero et al., 2014)) and the signal sequence is purported to be cleaved by a specialised signal peptidase, SipW (Stover & Driks, 1999a). Here we demonstrate that the first 43 amino acids are not needed for activity, as the TapA signal sequence can be replaced with the 28 amino acid TasA signal sequence, which is cleaved by SipW (Stover & Driks, 1999a, Stover & Driks, 1999b). When the chimeric construct, P_IPTG_-*tasAss-tapA*_*44-253*_, was expressed in the *tapA* deletion strain biofilm formation was fully reinstated (Fig. 3A). Therefore, the influence of amino acids 1-43 of TapA on protein function were excluded from further analysis.

**Figure 3:**
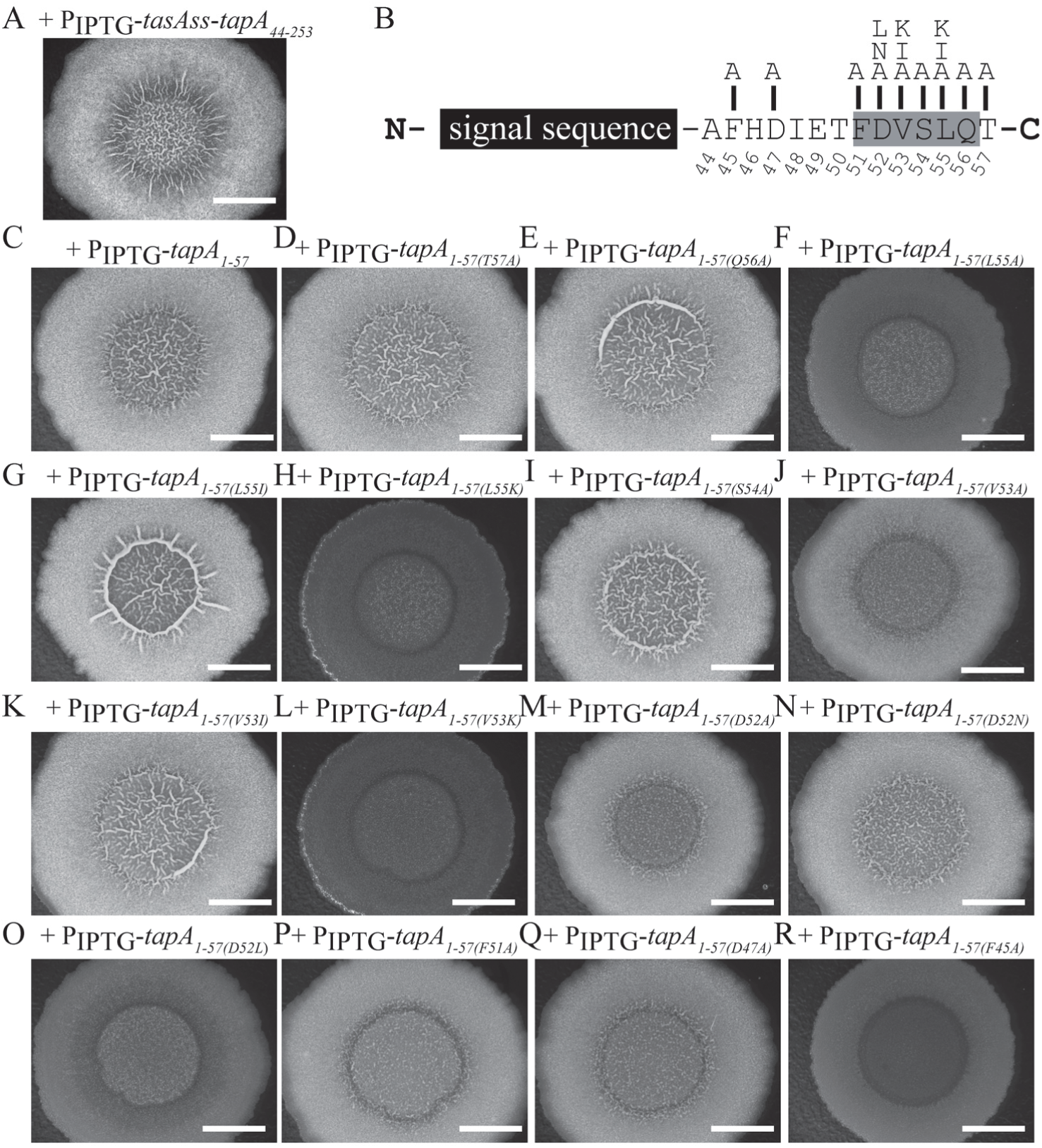
Identification of critical amino acids in the *tapA* coding region. (**A**) Biofilms formed by NRS6536 the *tasA* signal sequence replaces the native *tapA* signal sequence; (**B**) Schematic of the amino acid substitutions tested in **(C-R**); Biofilms formed by (**C**) +P_IPTG_-*tapA*_*1-57*_ (NRS6041; (**D**) +P_IPTG_-*tapA*_*1-57*_ T57A (NRS6384); (**E**) +P_IPTG_-*tapA*_*1-57*_ Q56A (NRS6385); (**F**) +P_IPTG_-*tapA*_*1-57*_ L55A (NRS6472); (**G**) +P_IPTG_-*tapA*_*1-*_ *57* L55I (NRS6386); (**H**) +P_IPTG_-*tapA*_*1-57*_ L55K (NRS6387); (**I**) +P_IPTG_-*tapA*_*1-57*_ S54A (NRS6388); (**J**) +P_IPTG_-*tapA*_*1-*_*57* V53A (NRS6473); (**K**) +P_IPTG_-*tapA*_*1-57*_ V53I (NRS6389); (**L**) +P_IPTG_-*tapA*_*1-57*_ V53K (NRS6502); (**M**) +P_IPTG_-*tapA*_*1-57*_ D52A (NRS6476); (**N**) +P_IPTG_-*tapA*_*1-57*_ D52N (NRS6516); (**O**) +P_IPTG_-*tapA*_*1-57*_ D52L (NRS6477); (**P**) +P_IPTG_-*tapA*_*1-57*_ F51A (NRS6390); (**Q**) +P_IPTG_-*tapA*_*1-57*_ D47A(NRS6475); and (**R**) +P_IPTG_-*tapA*_*1-57*_ F45A (NRS6474). Biofilms were grown at 30°C for 48 hours in the presence of 25 µM IPTG. n= 3. The scale bars represent 1 cm.

Next, site-directed mutagenesis was used to generate a series of constructs containing systematic substitutions in the *tapA*_*44-57*_ coding region in the context of the minimal TapA_1-57_ construct (Fig. 3B). The variant constructs were introduced into the *tapA* deletion strain and the ability of the variant TapA forms to restore biofilm formation assessed (Fig. 3C–3R). We first tested if the properties of the amino acid in position 57 were critical for activity. This was important to determine, as provision of the *tapA* coding region for amino acids 1-56 was unable to support biofilm recovery (Fig. 2C), whereas that encoding TapA_1-57_ was biologically active (Fig. 2C and 3C). A variant of TapA_1-57,_ where threonine 57 was replaced with alanine, had biological activity and therefore indicated that the chemical properties of amino acid 57 was less important than its presence (Fig. 3D). Therefore we concluded that the length of the TapA_1-57_ variant form is likely to be the important feature driving activity of TapA_1-57_.

We established that glutamine could substitute for alanine at amino acid position 56 (Fig. 3E). In contrast, replacement of leucine 55 with alanine generated a variant that was unable to support biofilm recovery in the *tapA* deletion strain (Fig. 3F). The importance of the hydrophobic nature of leucine 55 was supported as substitution with isoleucine, an amino acid with similar properties, was able to maintain the biological activity of TapA_1-57_ (Fig. 3G), while replacement with lysine, which mimics the size and shape of the leucine side chain but not the hydrophobicity, was not (Fig. 3H). We went on to show that the side chain of serine 54 was not essential for activity as TapA_1-57_ S54A fulfilled a role that was indistinguishable from the construct encoding TapA_1-57_ (Fig. 3I). Moving one amino acid along to valine 53, while both TapA_1-57_ V53A and TapA_1-57_V53K were unable to support formation of a rugose biofilm (Fig. 3J and 3L), the hydrophobicity of the side chain was essential with TapA_1-57_ V53I showing biological activity (Fig. 3K).

The replacement of aspartic acid at position 52 with either alanine (Fig. 3M) or asparagine (Fig. 3N) conferred (partial) structuring to the biofilm. In contrast TapA_1-57_ D52L had limited activity which demonstrates that replacement of an acidic amino acid with a large hydrophobic amino acid at position 52 is detrimental to TapA activity (Fig 3O). At position 51 changing phenylalanine to alanine resulted in biofilms with partial structure, indicating that TapA-_1-57_ F51A has limited function (Fig 3P). Finally two further features of the amino acid sequence stood out: 1) a conserved aspartic acid at position 47; and 2) an invariant phenylalanine at position 45. Constructs that resulted in individual substitution of these amino acids to alanine were generated. TapA_1-57_ D47A revealed only a partial ability to recover biofilm morphology (Fig. 3Q), while TapA_1-57_ F45A exerted no biological activity (Fig. 3L). In summary through this analysis we have identified amino acids F45, D52, V53 and L55 to be important for TapA function. These data underscore the importance of this stretch of amino acids to TapA function, however further analysis will be required to elucidate the role they play in facilitating TapA activity.

### TapA is detected *in vivo* at a low molecular mass

As the truncated variant of TapA covering amino acids 1-57 (inclusive) is sufficient for bioactivity, we wondered if TapA was processed to a smaller form *in vivo* and, if so, whether processing was a prerequisite for TapA function. To probe the molecular weight of TapA in the biofilm we first generated a custom TapA antibody using recombinant *B. subtilis* TapA_34-253_ as the antigen: a region that covers the mature secreted TapA protein. When the ability of the new antibody to bind to recombinant TapA_44-253_ protein was tested, a single band was detected at ~30 kDa (Fig. 4A), the apparent molecular mass where recombinant TapA_44-253_ is observed by SDS-PAGE. In contrast, immunoblot analysis of proteins extracted from the wild-type NCIB3610 sample revealed multiple distinct bands with apparent molecular masses of approximately 30 kDa, 18 kDa, and 16 kDa using αTapA antibodies. These bands were absent from the protein sample derived from the *tapA* mutant (Fig. 4B). The immunoreactive bands returned in an IPTG-dependent manner when proteins from the *tapA* complementation strain were analysed by immunoblot (Fig. 4B), indicating that they are dependent on TapA.

**Figure 4:**
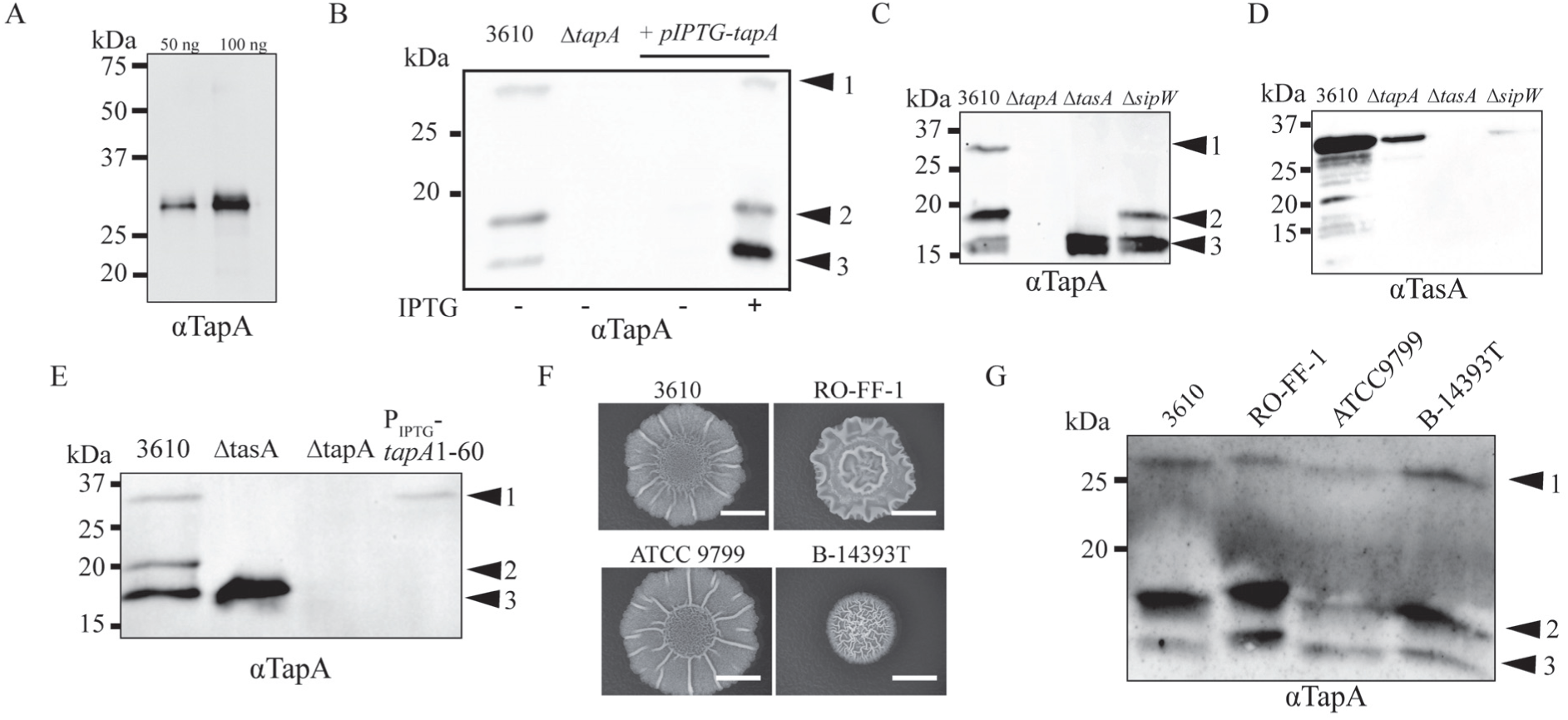
TapA is processed *in vivo*. (**A**) Immunoblot analysis of 50 and 100 ng of recombinant TapA_44-253_ using αTapA antibodies; (**B**) Immunoblot analysis of proteins extracted from biofilms formed by NCIB3610, Δ*tapA* (NRS3936), +P_IPTG_-*tapA* (NRS5045) (in the absence or presence of 25 µM IPTG as indicated) using αTapA antibodies. n= 2; (**C, D**) Immunoblot analysis of proteins extracted from biofilms formed by NCIB3610, Δ*tapA* (NRS3936), Δ*tasA* (NRS5267) and Δ*sipW* (NRS5488) strains using (**C**) αTapA and (**D**) αTasA antibodies. n= 2; (**E**) Immunoblot analysis of proteins extracted from biofilms formed in the presence of 25 µM IPTG by NCIB3610, Δ*tasA* (NRS5267), Δ*tapA* (NRS3936), +P_IPTG_-*tapA*_*1-60*_ (NRS6044) using αTapA antibodies. n= 2; (**F**) Biofilms formed by *B. subtilis* isolates NCIB3610, RO-FF-1, ATCC 9799, and B-14393T after growth at 30°C for 48 hours. Biofilm images are representative of at least 3 independent replicates. The scale bars represent 1 cm; (**G**) Immunoblot analysis of proteins extracted from biofilms formed by NCIB3610, RO-FF-1, ATCC 9799 and B-14393T using αTapA antibodies. n= 3.

Given the unexpected banding pattern observed using the proteins extracted from the wild type biofilm, further confirmation of the specificity of the αTapA antibody towards TapA was warranted. Considering the functional connections between TapA, TasA and SipW, proteins extracted from *tasA* and *sipW* deletion strains were examined by immunoblot using the αTapA antibody (Fig. 4C). For the *tasA* sample, a different banding profile to that of the wild-type was uncovered, with two of the three bands detected in the wild-type sample being absent. The third band, with an apparent molecular mass of 16 kDa, was still visible (Fig. 4C). The protein sample extracted from the *sipW* mutant displayed the lower 18 kDa and 16 kDa bands, however, as with the *tasA* sample, the 30 kDa band was again absent (Fig. 4C).

We also examined the protein samples using a TasA specific antibody (Erskine et al., 2018b) (Fig. 4D). We detected a dominant band at ~30 kDa for the wild-type strain, that was absent in the *tasA* sample and greatly reduced in the *tapA* and *sipW* strains (Fig. 4D). The reduction in the level of TasA is consistent with TapA and SipW influencing the amount of stable, secreted TasA (Romero et al., 2011). These findings also raised the possibility that the 30 kDa band detected using the αTapA antibody is actually TasA, not TapA: TasA is a dominant biofilm matrix protein that can be detected after coomassie staining of proteins analysed by SDS-PAGE (Erskine et al., 2018b, Branda et al., 2006). Consistent with the upper 30 kDa band being TasA, when proteins were extracted from a rugose biofilm where the truncated form of TapA (TapA_1-60_), which is capable of recovering biofilm formation, was expressed, the αTapA antibodies detected only the ~30 kDa band (Fig. 4E). We therefore conclude that the 16 kDa and 18 kDa bands are specific to TapA and the bands at 30 kDa and 18 kDa are proteins that are detected by the αTapA antibody when both TapA and TasA are present. The 30kDa band is likely to be TasA and the 16 kDa and 18 kDa bands processed forms of TapA. Overall the banding profile supports the conclusion that processing of TapA to smaller molecular weight forms occurs *in vivo*.

We next determined if low molecular mass TapA bands were detected in protein samples extracted from biofilms formed by other *B. subtilis* isolates. To do this we grew biofilms of *B. subtilis* isolates RO-FF-1, ATCC 9799 and B-14393T, alongside NCIB3610. After 48 hours incubation each of the isolates had formed a rugose biofilm (Fig. 4F). The mature biofilms were used to extract proteins corresponding to the total biofilm content and immunoblot analysis, using the αTapA antibody, clearly revealed that the TapA banding pattern was replicated in each of the distinct isolates (Fig. 4G). The consistent detection of TapA at a low molecular mass indicates that *in vivo* TapA may be subjected to processing in a controlled manner.

### TapA is cleaved by extracellular proteases

The detection of αTapA immunoreactive bands at low molecular weight raised the hypothesis that after secretion TapA is proteolytically cleaved in the extracellular environment, and that cleavage of TapA could release a functionally active form of the protein. *B. subtilis* encodes eight major secreted extracellular proteases that might fulfil this role named Bpr, Vpr, NprB, Mpr, Epr, AprE, NprE and WprA (Zhu & Stulke, 2018). To begin to explore this hypothesis, the susceptibility of recombinant TapA_44-253_ to proteolytic cleavage was first tested by incubating recombinant TapA_44-253_ with spent culture supernatant from strain NCIB3610, which contains the extracellular proteases secreted after growth to stationary phase. A control reaction for this experiment where the spent culture supernatant was heat treated for 10 min at 100°C to denature the native exoproteases was also implemented. 120 µg ofrecombinant TapA_44-253_ was added to the culture supernatant and incubated for 8 hrs at 37°C. When the samples were analysed by SDS-PAGE the full length protein was no longer detectable when incubated with untreated culture supernatant, but the protein remained at full size when incubated with the heat treated culture supernatant (Fig. S2A). Taken together with data indicating that TapA is a secreted protein, these findings suggest that the extracellular proteases can degrade TapA *in vivo*.

### Proteolysis of TapA is not needed for biofilm formation

To test the potential impact of blocking proteolytic cleavage of TapA on biofilm formation, and thereby establish if proteolysis of TapA was a requirement for activity, we constructed a strain that lacked seven of the secreted exoproteases (hereafter KO7: NRS6362) and another which lacked all eight secreted exoproteases (hereafter KO8: NRS5645). Neither of the strains generated showed any evidence of proteolytic activity when grown on LB agar supplemented with 1.5% (w/v) milk, similar to the *degU* deletion strain which was used as a negative control (Fig. S2B) (Msadek *et al.*, 1990).

We first tested the stability of recombinant TapA protein in spent culture supernatant isolated from the KO7 and KO8 strains. After incubation there was limited evidence of proteolysis of recombinant TapA_44-253_ when incubated with KO7 supernatant and no observable degradation of TapA_44-_ 253 when it was incubated with KO8 supernatant (Fig. S2A). Consistent with these findings, immunoblot analysis of proteins extracted after growth for 48 hours under biofilm formation conditions revealed that *in vivo* processing of TapA to the αTapA specific 16 kDa form was impeded in both the KO7 and KO8 strains. The 18 kDa and 30 kDa bands were still detected but additional bands of intermediate sizes were also apparent (Fig. 5A). We postulate that the additional bands in the KO7 and KO8 strains represent differentially cleaved TapA fragments.

**Figure 5:**
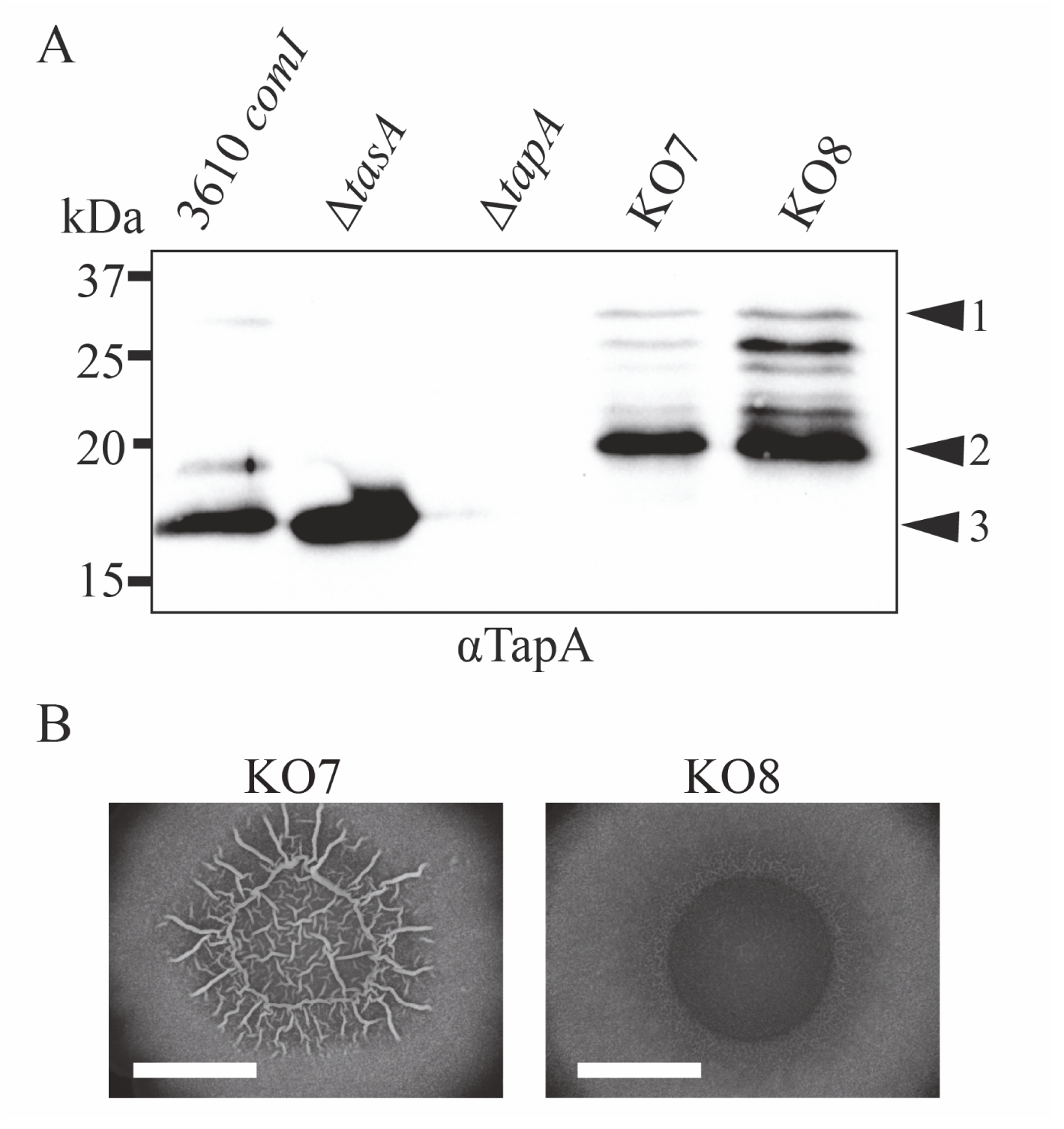
Proteolytic cleavage of TapA is not a prerequisite for biofilm formation. (**A**) Immunoblot analysis of proteins extracted from biofilms formed by NCIB3610, Δ*tasA* (NRS5267), Δ*tapA* (NRS3936), KO7 (NRS6362) and KO8 (NRS5645) using αTapA antibodies; n = 2. (**B**) Biofilms formed by *B. subtilis* isolates KO7 (NRS6362) and KO8 (NRS5645) after growth at 30°C for 48 hours. Biofilm images are representative of at least 3 independent replicates. The scale bars represent 1 cm.

Despite the *in vivo* block in TapA proteolysis, no obvious impact on the structure of the mature biofilm was detected for the KO7 strain (Fig. 5B). In contrast, the KO8 strain displayed a defect in biofilm formation, lacking the robust structuring typical of the wild-type strain (Fig. 5B). The single *wprA* deletion strain, the last gene to be deleted in the KO8 strain, showed no alternation in biofilm formation (Fig. 2SD). Therefore the lack of wrinkles in the biofilm formed by the KO8 strain was not specific to the absence of WprA. The lack of rugosity is not likely to be a consequence of a generalised growth defect as both the wild type and KO8 strain had comparable growth rates and yields in MSgg under shaking culture conditions (Fig. S2C).

To determine if the reduction in rugosity displayed by the KO8 strain was due to a lack of TapA processing, we introduced the minimal functional unit coding region, TapA_1-57_, at the *amyE* location and assessed biofilm formation. We hypothesized that if processing of TapA was essential for function introduction of TapA_1-57_ should reinstate biofilm architecture. In opposition to this hypothesis, it was found that production of TapA_1-57_ was unable to re-instate rugosity of KO8 biofilms such that it looked like the wild-type strain (NRS6960) (Fig. S2E). Taken together with the immunoblot data demonstrating the same TapA banding pattern in the KO7 and KO8 strains, these findings indicate that proteolytic cleavage of TapA by extracellular proteases is not a prerequisite to generate a functionally active form of TapA.

### TapA is not a chaperone protecting TasA from exoprotease activity

The exact role of TapA during biofilm formation is elusive having been linked with TasA stability (Romero et al., 2011), TasA-fibre formation and attachment (Romero et al., 2010, Romero et al., 2011) and forming fibres itself (El Mammeri et al., 2019). The construction of the exoprotease minus strain allowed us to test the possibility that it functions as a chaperone. The formation of exoprotease-resistant fibres by TasA is linked with TapA *in vivo*, with the levels of TasA being reduced in the *tapA* mutant strain (Fig 4D). Moreover, TasA is prone to degradation by the exoproteases, when it is not in fibre form (Fig. S3A and S3B) (Erskine et al., 2018b). Therefore one hypothesis is that TapA is a chaperone that shields monomeric TasA from proteolytic degradation in the extracellular environment, thereby allowing TasA fibres to form. If correct, we predicted that TapA would not be needed for TasA fibre formation, and consequentially biofilm formation, in a strain lacking the extracellular proteases. Consistent with this hypothesis, recombinant TasA, that is restricted to a monomeric form by the addition of a single amino acid at the N-terminus (Erskine et al., 2018b), is only prone to degradation by the extracellular proteases found in the wild type culture supernatant. The recombinant protein is not degraded when incubated in culture supernatant isolated after growth of the exoprotease-free KO8 strain (Fig. S3A) (Erskine et al., 2018b). To determine if *tapA* would be needed for biofilm formation in the absence of extracellular proteases, we deleted *tapA* from the KO7 and KO8 strains and examined biofilm formation. The biofilms formed by strains KO7 Δ*tapA* (NRS6293) and KO8 Δ*tapA* (NRS5646) were examined and contrary to our hypothesis both the KO7 Δ*tapA* and KO8 Δ*tapA* formed biofilms with the flat, featureless appearance of the Δ*tapA* strain (Fig 6A and 6B). In each case the biofilm defect was specific to the loss of *tapA* as biofilm formation was reinstated when the full length *tapA* coding region was expressed from the *amyE* location (Fig. 6A and 6B). Additionally we were able to demonstrate that TapA_1-57_ was still functional in this system: we found that it could restore the colony morphology of KO8 Δ*tapA* biofilms (NRS6961) (Fig. S2F) to similar levels as displayed by the KO8 strain (NRS5645) (Fig. S2E). Together these data indicate that *tapA* does not become dispensable during biofilm formation in the absence of extracellular proteases and therefore is unlikely to serve as a chaperone to protect monomeric TasA from degradation during fibre formation.

**Figure 6:**
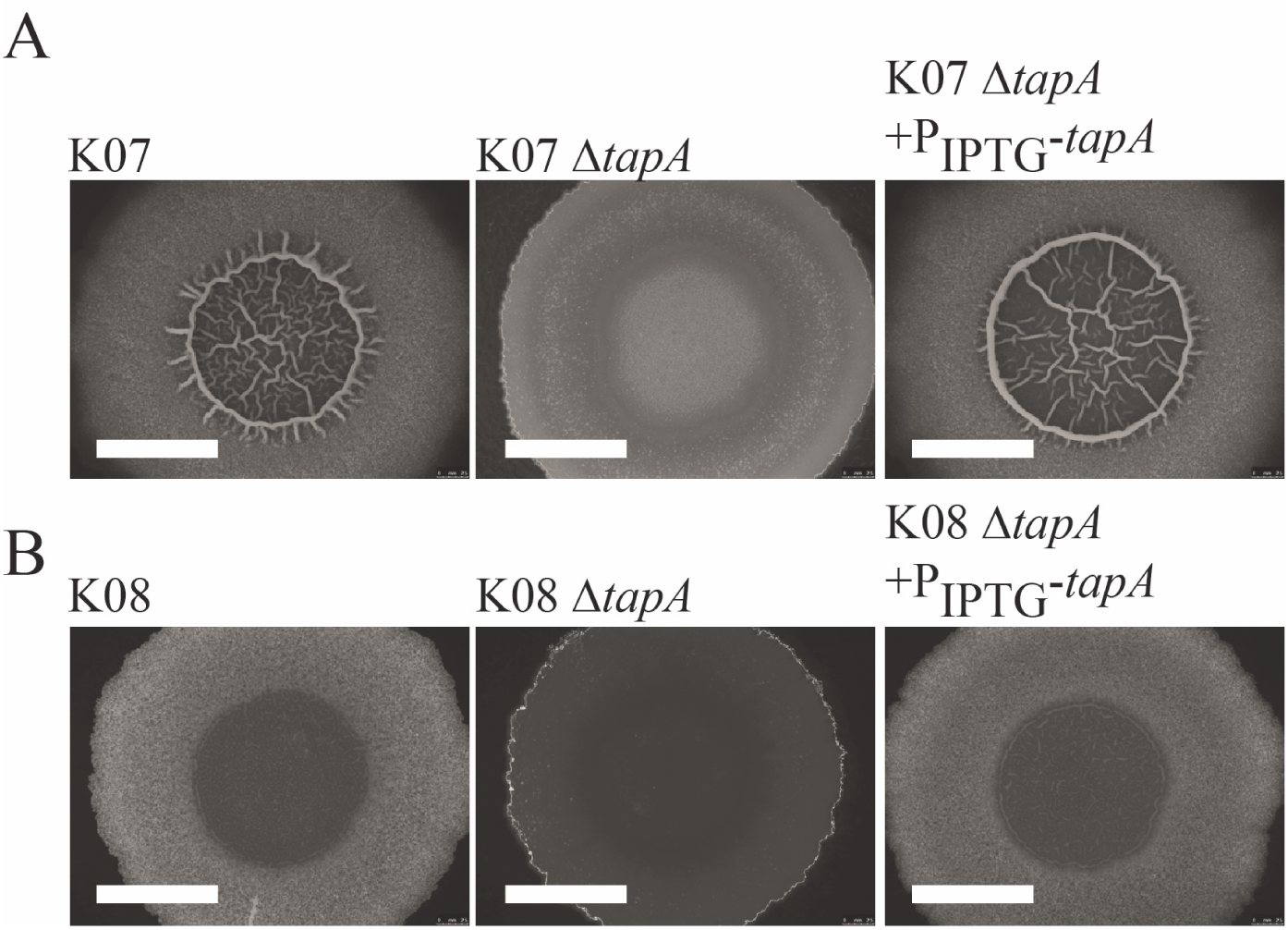
The *tapA* coding region is needed for biofilm formation in the absence of extracellular proteases. Biofilms formed by *B. subtilis* isolates (**A**) KO7 (NRS6362), KO7 Δ*tapA* (NRS6293), KO7 Δ*tapA* P_IPTG_-*tapA*-*lacI* (NRS6295); (**B**) KO8 (NRS5645), KO8 Δ*tapA* (NRS5646), KO8 Δ*tapA* P_IPTG_-*tapA*-*lacI* (NRS5647); after growth at 30°C for 48 hours. Biofilm images are representative of at least 3 independent replicates. The scale bars represent 1 cm.

## Discussion

An extracellular matrix comprised of polymers is critical to structure sessile communities of bacterial cells called biofilms. Biofilm matrix production by *Bacillus subtilis* depends on the protein TapA, which has a role in promoting TasA stability and fibre formation *in vivo*. Here we show that orthologous *tapA* genes, originating from other *Bacillus* species, are functional in *B. subtilis* NCIB3610 strain. This is in agreement with previous work which found that *tapA* from *B. amyliquofacienes* can substitute for the *B. subtilis tapA* coding region (Romero et al., 2014). The *B. pumilis* TapA protein is 61 amino acids shorter at the C-terminus, compared with *B. subtilis* TapA, indicating that the extreme C-terminal amino acids are dispensable for rugose biofilm formation. Consistent with this, when a truncated form of the *B. subtilis tapA* open reading frame was used to encode a variant form of TapA that lacked the C-terminus it was functional. Further investigations presented here determined that the minimal functional unit of TapA consists of amino acids TapA _1-57_. This was unexpected as the region of TapA with the highest conservation amongst the orthologues (amino acids 58-253 inclusive) was not needed to restore rugose biofilms to the *tapA* deletion strain. We have also shown that the predicted TapA signal sequence, which consists of amino acids 1-43, can be replaced with the predicted TasA signal sequence. This therefore leaves a functional component of TapA of only 14 amino acids (amino acids 44-57 inclusive). We cannot, however, rule out the possibility that amino acids 58-253 of TapA serve a distinct function beyond rugose biofilm formation. Indeed, recent work has suggested that the C-terminus of TapA is a disordered protein while the N-terminus is a structured domain, with the 2 domains interacting with lipid vesicles in a cooperative manner (Abbasi et al., 2019).

Further to the discovery of the minimal functional form of TapA, we uncovered that TapA was processed *in vivo* to lower molecular weight forms in the biofilm. We also observed processed forms of TapA in the Δ*sipW* mutant which is suggestive of the conclusion SipW is not essential for TapA secretion. It could be that another signal peptidase(s) can substitute for the loss of *sipW* in terms of TapA secretion or that TapA is retained associated with the cell membrane *via* its signal peptide when it is processed. The cleavage of TapA to lower molecular weight forms is conserved amongst other *B. subtilis* isolates and depends on extracellular proteases. We speculated that TapA processing might be needed to generate an active variant of TapA and tested the impact of deleting the genes encoding the extracellular proteases on biofilm formation. While we found that TapA processing to lower molecular weight forms was altered both in a strain lacking Bpr, Vpr, NprB, Mpr, Epr, AprE, and NprE (KO7) and a second where WprA was also deleted (KO8), in contradiction to our hypothesis, although there was a partial defect in biofilm formation exhibited by the KO8 strain, the biofilm formed by KO7 resembled wild-type morphology. As analysis of the TapA protein from these biofilms showed a similar banding pattern using immunoblot in both the KO8 and KO7 strains, we conclude that the KO8 biofilm defect is likely to be due to pleiotropic effects of exoprotease gene deletion, rather than being a specific impact of altered TapA processing. These findings raise the possibility that TapA is degraded (recycled) after it has fulfilled its role in biofilm formation or support the notion that TapA plays a second role in *B. subtilis* physiology that is yet to be elucidated.

Having ruled out that the defect in biofilm formation observed for the exoprotease-free KO8 strains is not because of aberrant TapA processing, the reason why the loss of eight extracellular proteases impacts biofilm rugosity remains unknown. It is possible that the lack of extracellular proteases could impact the abundance of quorum sensing peptides in the extracellular environment. This is in line with evidence demonstrating that exoproteases control both the production and degradation of the quorum sensing signalling molecule ComX (Spacapan *et al.*, 2018). An accumulation of quorum sensing peptides in the biofilm microenvironment may have a global impact on the physiology of the KO8 strain due to an alteration in cell signalling pathways (Kalamara *et al.*, 2018, Miller & Bassler, 2001).

The generation of a strain that lacked the known extracellular proteases allowed us to test the hypothesis that TapA functions as a chaperone during TasA fibre formation. This hypothesis was raised as while fibrous TasA (fTasA) is resistant to proteolysis when TasA is in its monomeric form (mTasA) it is sensitive to the action of proteases. When this knowledge was coupled with reduction of TasA levels in a *tapA* mutant, it was hypothesised that TapA could serve as a chaperone interacting with mTasA and preventing degradation to facilitate fibre formation. In this model, when the exoproteases are absent, TapA would not be needed for wild-type biofilm formation. However, when the morphology of the KO7 Δ*tapA* and KO8 Δ*tapA* strains were analysed they both resembled the appearance of a Δ*tapA* mutant. These findings therefore indicate that TapA does not play a significant role in protecting TasA from proteolysis by the extracellular enzymes.

### Concluding Remarks

Through this analysis we have expanded our knowledge of the protein TapA which is needed for biofilm formation by *B. subtilis*. We have revealed that TapA_1-57_ is a key component of the functional form of TapA *in vivo*, allowing biofilm formation. The main body of the protein is entirely dispensable in this experimental set up. We have shown that TapA is processed *in vivo* but that processing by the self-produced exoproteases is not an essential step to generate a functional protein form. The exact mechanism by which TapA is needed to stimulate TasA stability *in vivo* requires more elucidation, but the amino acids that were found to be critical for function between amino acids 45-57 provide a starting point for analysis. Finally we have ruled out the possibility that *tapA* is not needed when extracellular proteases are not present. Therefore exactly how TapA functions to promote biofilm formation remains to be elucidated.

### Growth media and additives

Lysogeny broth (LB) was prepared using 10 g typtone, 10 g NaCl and 5 g yeast extract for 1 litre. LB agar was prepared by solidifying with 15 g of select agar for growth of *B. subtilis* and *E. coli*. For antibiotic selection with *B. subtilis* strains antibiotics were used at the following final concentrations: erythromycin (0.5 µg/mL), spectinomycin 100 µg/mL and MLS, erythromycin (0.5 µg/mL) together with lincomycin (12.5 µg/mL). For antibiotic selection of plasmids in *E. coli* ampicillin was used at a concentration of 100 µg/mL.

To make Minimal Salts glycerol glutamate (MSgg) plates 5 mM potassium phosphate, 100 mM MOPS at pH 7.0 with agar to a final concentration of 1.5% (w/v) was autoclaved and cooled to 55°C before being supplemented with 2 mM MgCl_2_, 700 µM CaCl_2_, 50 µM FeCl_3_, 50 µM MnCl_2_, 1 µM ZnCl_2_, 2 µM thiamine, 0.5% (v/v) glycerol, 0.5% (w/v) glutamic acid. For induction of gene expression from the P_spank_ promoter (P_IPTG_) isopropyl β-D-1-thiogalactopyranoside (IPTG) was included at a final concentration of 25 µM. For MSgg broth the same recipe was followed without the addition of agar.

### Plasmid construction

Plasmids (Table S2) were constructed by using standard methods using the primers detailed in Table S3.

### Strain construction

Strains used in this study are detailed in Table S1. Complementation alleles and antibiotic resistance cassette marked gene deletions were moved between strains using either SSP1 mediated phage transduction (Verhamme *et al.*, 2007) or genetic competence with genomic DNA. The following strains were used for amplification of coding regions: *B. subtilis* strain NCIB 3610 (GenBank Accession number: CP020102.1); *B. amyloliquefaciens* FZB42 (GenBank Accession number: CP000560.1); B. *paralicheniformis* (kindly provided by Dr. Nijland) and *B. pumilis* SAFR-032 (GenBank Accession number: CP000813.4).

The Δ*tapA* and Δ*sipW* deletion strains were generated by allelic exchange in a method similar to that previously published using the pMAD plasmid (Arnaud *et al.*, 2004). Briefly, a 395 bp upstream region was amplified by PCR with primers NSW1308 and NSW1332 and a 641 bp downstream region was amplified using primers NSW1333 and NSW1334. Both fragments were cloned into the pMini-MAD vector (Patrick & Kearns, 2008a) to generate plasmid pNW685. For the *sipW* deletion strain a 1,128 bp fragment containing 572 bp of sequence upstream of the *sipW* locus followed by a 36 bp open reading frame in place of *sipW*, with 572 bp of sequence downstream of *sipW* was synthesised (GenScript). The fragment was cloned into the pMini-MAD vector to generate plasmid pNW2021.

To construct the in frame deletions in all 8 genes encoding the secreted proteases *B. subtilis* NCIB3610 *comI* (Patrick & Kearns, 2008a), the BKE collection was utilised (Koo *et al.*, 2017b) where single gene deletions have been replaced with a cassette providing resistance to erythromycin. Genomic DNA was extracted from strains in the BKE collection and used to transform competent *B. subtilis* 3610 *comI* (Konkol *et al.*, 2013) before selection on LB erythromycin plates. The erythromycin cassette contains *lox* sites and was subsequently removed leaving a 150 base pair scar by the action of a Cre recombinase, which was expressed on the heat-sensitive plasmid pDR244. In cases in which transformation with genomic DNA proved unsuccessful then the mutation was introduced by phage transduction with SPP1 phage. All the strains were examined using PCR and DNA sequencing to ensure the specificity in the region deleted from the chromosome. The intermediate strains are fully detailed in Table S1. The in frame *tapA* deletion was introduced using plasmid pNW685 described above.

The variant *tapA* coding regions were introduced into *B. subtilis* chromosome at the *amyE* locus. Double recombination events were identified by assessing the production of α-amylase on LB growth medium supplemented with 1% (w/v) soluble starch.

### Growth analysis

Lawn plates were set up by suspending a single colony in 100 µL of LB medium and plating the suspension onto LB agar and incubated overnight at 25°C. After ~24 h the cells were washed from the plate in 5 mL of MSgg broth and the absorbance at 600 nm measured. The volume of cell suspension used to inoculate was calculated based on a desired starting OD_600nm_ of 0.1. Cultures were grown in 25 mL of MSgg broth in a 250 mL conical flask in a water bath set to 30°C with shaking at 200 rpm.

### *In vivo* analysis of exoprotease production

Detection of exoprotease production was conducted as previously described (Verhamme et al., 2007) using LB agar plates supplemented with 1.5% (w/v) dried milk powder. Strains were grown in 3 mL of LB broth at 37°C to an OD_600nm_ of ~ 1. The cultures were normalized and 10 µL of the prepared cell culture spotted onto the plate. The samples were then grown for 20 h at 37°C prior to photography.

### Biofilm growth and analysis

Biofilms were prepared by growing *B. subtilis* in 3 mL of LB broth at 37°C with aeration for ~3.5 hours. After which, 10 µL of the culture was spotted onto an MSgg agar plate that was incubated at 30°C for 48 hours. Imaging used a Leica MZ16 stereoscope (Leica Microsystems). IPTG was included in the growth medium to induce expression from the P_spank_ promoter as indicated: a concentration of 25 µM IPTG was used, unless otherwise stated.

### Protein extraction from biofilms

Biofilms were isolated from MSgg plates with a sterile loop and suspended in 250 µL of BugBuster solution (Millipore) using a syringe with a 23 × 1 gauge needle until dispersed. The samples were sonicated at an amplitude of 20% power for 5 seconds. Sonicated biofilms were incubated at 26°C for 20 min with shaking at 1,400 rpm before centrifugation for 10 min in a benchtop centrifuge at 13,000 rpm. The liquid phase was retained for further analysis by SDS-PAGE and/or immunoblot. Protein concentration of biofilm lysates was determined by measuring absorbance at 280 nm (NanoDrop spectrophotometer) or by using the DC protein assay (BIO-RAD) which is based on the Lowry assay (Lowry *et al.*, 1951).

### Protein purification

Recombinant *B. subtilis* TapA_44-253_, mTasA (Erskine et al., 2018b) and fTasA (Erskine et al., 2018b) proteins were produced and separated from a Glutathione S-transferase-tag with a tobacco etch virus (TEV) protease-cleavage site using the pGEX-6P-1 system. The pGEX-6P-1 plasmid carrying the gene encoding the protein was introduced into *E. coli* strain BL21 (DE3) pLysS. The cells were grown overnight in 5 mL LB broth and used to inoculate auto-induction media (Studier, 2005) supplemented with ampicillin (100 µg/mL) at a ratio of 1:1000 (vol:vol) in a total volume of 1 litre. The cultures were incubated at 30°C with shaking for approximately 6-7 h at which point the temperature was reduced to 18°C for overnight incubation. The cell culture was pelleted by centrifugation for 45 min at 5020 g and re-suspended in 25 mL purification buffer (Tris 25 mM and NaCl 250 mM (pH 7.6)) supplemented with Complete EDTA-free proteinase inhibitors mixture (Roche). Cell lysis was carried out by sonication at an amplitude of 20% for a total of 6 minutes. Unlysed cells and cell debris were removed by centrifugation at 27,000 x g for 20 min. The cleared lysate was mixed with 750 µL (per litre of culture) Glutathione Sepharose 4B (GE Healthcare) and gently agitated at 4°C for at least 3 h to allow binding of GST to the beads. The lysate/bead mixture was loaded onto a single-use, 25 mL gravity flow column (Bio-Rad). The beads were washed using 50 mL purification buffer, collected and incubated overnight at 4°C with agitation in 25 mL of purification buffer supplemented with 1 mM DTT and 0.5 mg of TEV protease to release the protein from the GST-tag. The solution containing TapA, TEV protease, free-GST and the beads was loaded onto the gravity flow column. This removed the used beads which stay on the column. The flow-through was added to 250 µL Ni-nitrilotriacetic acid agarose (Qiagen) slurry to remove the TEV protease and 750 µL glutathione sepharose 4B to remove the free GST-tag. The mixture was incubated at 4°C with agitation overnight, and then passed through a gravity flow column. The purified protein was concentrated using a Vivaspin 20 concentrator (with a MW cut-off of 5,000 or 10,000 Sartorius). The protein concentration was determined by measuring absorbance at 280 nm (NanoDrop spectrophotometer) and then analysed by separating ~30 µg by SDS-PAGE.

### Protein stability in culture supernatant

To collect spent culture supernatant the following process was used. Initially lawn plates were set up by suspending a single colony in 100 µL of LB medium and plating the suspension onto LB agar. Lawn plates were incubated overnight at 25°C. After ~24 h the cells were washed from the plate in 5 mL of MSgg broth and the absorbance at 600 nm measured. The volume of cell suspension used to inoculate was calculated based on a desired starting OD_600nm_ of 0.1. Cultures were grown in 25 mL of MSgg broth in a 250 mL conical flask in a water bath set to 30°C with shaking at 200 rpm. Early stationary phase cultures were harvested at an OD_600_ of approximately 4.0 and normalized. The harvested culture was then pelleted by centrifugation at 3220 x g for 10 min and the full volume of supernatant was filter-sterilized to remove bacterial cells. The supernatant protease activity assay was set up by mixing supernatant 1:1 (vol:vol) with purified TapA, mTasA or fTasA protein (to give a total volume of 40 µL) and incubating the mixture at 37°C for 8 h. This gave a protein concentration in the assay of 3 µg/µL, and a total recombinant protein amount of 120 µg. Heat-inactivation of the supernatant was carried out by incubating the supernatant samples at 100°C for 10 min prior to use. The integrity of the recombinant protein was then assessed by SDS-PAGE, stained with InstantBlue, Coomassie based stain. 28 µg of protein was loaded onto the gel, this was calculated based on the starting assay concentration of 3 µg/µL.

### Antibody Production

A custom antibody that could be used to detect TapA from *B. subtilis* was raised in a rabbit using purified recombinant TapA_34-253_ as the antigen (Eurogentec). The antibodies specific to recombinant TapA_34-253_ were purified from the serum using standard methods by the MRC Protein reagents and services team (https://mrcppureagents.dundee.ac.uk/our-services/custom-antibody-production).

### Immunoblot analysis

Samples to be analysed by immunoblotting were separated by SDS-PAGE. The proteins were transferred to hydrophobic polyvinylidene difluoride (PVDF) membranes (Immobilon-P (Millipore)) by electroblotting. The membranes were first incubated with a 5% (w/v) semi-skimmed dry milk solution in TBS-tween 0.2% (v/v) for at least 1 hour at room temperature or overnight at 4°C to reduce non-specific binding. After which the membrane was incubated with the primary antibody overnight at 4°C (dilution as follows: 1:5000 αTapA, 1:25,000 αTasA). After incubation with the primary antibody, the membrane was washed 3 times with TBS-tween 0.2% (v/v) to remove unbound primary antibody and incubated with the species-specific secondary HRP-conjugated antibody (Goat α Rabbit, dilution 1:5000) for 1 hour at room temperature in TBS-tween 0.2% (v/v). The wash steps were repeated before development was induced with Enhanced Chemi-Luminescence reagents (ECL; BioRad Clarity). An electronic imaged was captured using the GeneGnome (SynGene) system.

### Bioinformatics analysis

Protein sequences of TapA homologues were aligned using Clustal Omega using default settings (Sievers *et al.*, 2011). Percentage identity between TapA orthologues was calculated with reference to *B. subtilis* TapA using the pairwise alignment function on the Jalview 2 workbench (Waterhouse *et al.*, 2009). For the prediction of signal peptides for all TapA variants then the SignalP 4.1 server was used and set to the organism group “Gram-positive” (Petersen et al., 2011).

## Acknowledgements

Work was supported by the Biotechnology and Biological Sciences Research Council [BB/L006804/1; BB/L006979/1; BB/M013774/1; BB/N022254/1]. CE was supported by the Wellcome Institutional Strategic Support Fund (Award no. 097818/Z/11). We are grateful to Dr. Hobley for the *tapA* deletion strain, Dr. Elliot Erskine for the *sipW* deletion strain, Dr. Laura D’Ignazio for plasmid pNW1600 and Rachel Gillespie for construction of a number of strains and plasmids.

