## Supplemental information for "The majority of the matrix protein TapA is dispensable for biofilm formation by *Bacillus subtilis*"

**Synthesised genetic constructs**

**Δ*sipW* construct (GenScript)** (cloning sites EcoRI-SalI) sub-cloned to create pNW2021

gaattcTAAAGACTTTCAGCATACAGATAAAAACTGCCATTATGATAAACGCTGGGATCAAAGTGATTTGCACATATCAGATCAA
ACGGATACGAAAGGCACTGTATGCTCACCTTTCGCCTTATTTGCTGTGCTCGAAAATACAGGTGAGAAACTTAAGAAATCAAA
GTGGAAGTGGGAGCTTCATAAGCTTGAAAATGCCCGCAAACCGTTAAAGGATGGGAACGTGATCGAAAAAGGATTTGTCTCC
AATCAAATCGGCGATTCACTTTATAAAATTGAGACCAAGAAAAAAATGAAACCCGGCATTTATGCATTTAAAGTATATAAACCG
GCAGGCTACCCGGCAAACGGCAGTACATTTGAGTGGTCGGAGCCTATGAGGCTTGCAAAATGCGATGAAAAACCGACAGTCC
CTAAAAAAGAAACAAAGTCGGACGTCAAAAAGGAGAATGAAACAACACAAAAAGATATACCGGAAAAAACAATGAAAGAAG
AAACATCTCAAGAAGCTGTAACCAAAGAAAAAGAAACTCAATCAGACCAGAAGGAAAGCGGGGAAGAGGATGAAAAAAGCA
**ATG**AAGCTGATCAGTAATATTTTATACTCTACT**TAA**CTTCAGTTGTAAACCTGGCAACAGGTTTCGATATAAAATCATTCAATAA
AAGGGGAGCTTACCATGGGTATGAAAAAGAAATTGAGTTTAGGAGTTGCTTCTGCAGCACTAGGATTAGCTTTAGTTGGAGG
AGGAACATGGGCAGCATTTAACGACATTAAATCAAAGGATGCTACTTTTGCATCAGGTACGCTTGATTTATCTGCTAAAGAGA
ATTCAGCGAGTGTGAACTTATCAAATCTAAAGCCGGGAGATAAGTTGACAAAGGATTTCCAATTTGAAAATAACGGATCACTT
GCGATCAAAGAAGTTCTAATGGCGCTTAATTATGGAGATTTTAAAGCAAACGGCGGCAGCAATACATCTCCAGAAGATTTCCT
CAGCCAGTTTGAAGTGACATTGTTGACAGTTGGAAAAGAGGGCGGCAATGGCTACCCGAAAAACATTATTTTAGATGATGCG
AACCTTAAAGACTTGTATTTGATGTCTGCTAAAAATGATGCAGCGGgtcgac

***tapA***$_{RBS}$-***tasA***$_{1\text{-}27}$-***tapA***$_{44\text{-}253}$ **(DCBiosciences)** (cloning sites SalI-SphI) sub-cloned to create pNW1885

gtcgacTTTTACAGGAGGTAAGATATGGGTATGAAAAAGAAATTGAGTTTAGGAGTTGCTTCTGCAGCACTAGGATTAGCTTTA
GTTGGAGGAGGAACATGGGCAGCTTTTCATGATATTGAAACATTTGATGTCTCACTTCAAACGTGTAAAGACTTTCAGCATACA
GATAAAAACTGCCATTATGATAAACGCTGGGATCAAAGTGATTTGCACATATCAGATCAAACGGATACGAAAGGCACTGTATG
CTCACCTTTCGCCTTATTTGCTGTGCTCGAAAATACAGGTGAGAAACTTAAGAAATCAAAGTGGAAGTGGGAGCTTCATAAGCT
TGAAAATGCCCGCAAACCGTTAAAGGATGGGAACGTGATCGAAAAAGGATTTGTCTCCAATCAAATCGGCGATTCACTTTATA
AAATTGAGACCAAGAAAAAAATGAAACCCGGCATTTATGCATTTAAAGTATATAAACCGGCAGGCTACCCGGCAAACGGCAGT
ACATTTGAGTGGTCGGAGCCTATGAGGCTTGCAAAATGCGATGAAAAACCGACAGTCCCTAAAAAAGAAACAAAGTCGGACG
TCAAAAAGGAGAATGAAACAACACAAAAAGATATACCGGAAAAAACAATGAAAGAAGAAACATCTCAAGAAGCTGTAACCAA
AGAAAAAGAAACTCAATCAGACCAGAAGGAAAGCGGGGAAGAGGATGAAAAAAGCAATGAAGCTGATCAGTAAgcatgc

**Table S1 Strains used in this study**

| Strain | Relevant genotype/description [a] | Source/construction [b] |
|---|---|---|
| 168 | *trpC2* | BGSC |
| BL21 | F–*ompT hsdSB (rB–), (mB–)gal dcm* (DE3) | (Studier & Moffatt, 1986) |
| MC1061 | *F' lacIQ lacZM15 Tn10 (tet)* | |
| NCIB3610 | *prototroph* | BGSC |
| RO-FF-1 | *B. subtilis* wild isolate (stocked as NRS1144) | (Roberts & Cohan, 1995) |
| NRS1314 | NCIB3610 *degU::pBL201 (cml)* | (Verhamme et al., 2007) |
| NRS2661 | *E. coli* BL21 pGEX-6P-1+TEV-site+*tapA*$_{B\_sub44-253aa}$ | pNW1600→*E. coli* BL21 |
| NRS5028 | *E. coli* BL21 pGEX-6P-1+TEV-site+*tasA*$_{B\_sub28-261aa}$ | pNW1437→*E. coli* BL21 |
| NRS5253 | *E. coli* BL21 pGEX-6P-1+TEV-site+ser-*tasA*$_{B\_sub28-261aa}$ | pNW1080→*E. coli* BL21 |
| NRS3789 | 168 pMiniMAD Δ*tapA* | pNW685→168 |
| NRS3936 | NCIB3610 *ΔtapA* | Laura Hobley (This work) |
| NRS5487 | NCIB3610 *ΔsipW* | pNW2021→168<br>(Elliot Erskine) |
| NRS5488 | 168 pMiniMAD *ΔsipW* | SPP1 NRS5487→ NCIB3610<br>(Elliot Erskine) |
| NRS5034 | *E. coli* BL21 pGEX-6P-1+TEV-site+*tapA*$_{B\_sub34-253aa}$ | pNW1441→*E. coli* BL21 |
| NRS5041 | 168 *amyE::P-spank-tapA*$_{B\_sub*}$*-lacI (spc)* | pNW1438→168 |
| NRS5042 | 168 *amyE::P-spank-tapA*$_{B\_pum**}$*-lacI (spc)* | pNW1439→168 |
| NRS5043 | 168 *amyE::P-spank-tapA*$_{B\_amy***}$*-lacI (spc)* | pNW1440→168 |
| NRS5045 | NCIB3610 *ΔtapA + amyE::P-spank-tapA*$_{B\_sub}$*-lacI (spc)* | SPP1 NRS5041→NRS3936<br>(Rachel Gillespie) |
| NRS5046 | NCIB3610 *ΔtapA + amyE::P-spank-tapA*$_{B\_pum}$*-lacI (spc)* | SPP1 NRS5042→NRS3936<br>(Rachel Gillespie) |
| NRS5047 | NCIB3610 *ΔtapA + amyE::P-spank-tapA*$_{B\_amy}$*-lacI (spc)* | SPP1 NRS5043→NRS3936<br>(Rachel Gillespie) |
| NRS5048 | 168 pMiniMAD Δ*tasA* | pNW1448→168 |
| ATCC 9799 | *B. subtilis* wild isolate (stocked as NRS5142) | (Duthie, 1944) BGSC:3A14 |
| B-14393T | *B. subtilis* wild isolate (stocked as NRS5145) | (Priest *et al.*, 1988)<br>BGSC:10A5 |
| NRS5267 | NCIB3610 *ΔtasA* | (Erskine et al., 2018b) |
| NRS5741 | 168 *amyE::P-spank-tapA*$_{B\_para****}$*-lacI (spc)* | pNW1800→168 |
| NRS5742 | 168 *amyE::P-spank-tapA*$_{B\_sub\_1-193}$*-lacI (spc)* | pNW1801→168 |
| NRS5743 | NCIB3610 *ΔtapA + amyE::P-spank-tapA*$_{B\_para}$*-lacI (spc)* | SPP1 NRS5741→NRS3936 |
| NRS5744 | NCIB3610 *ΔtapA + amyE::P-spank-tapA*$_{Bs\_1-193}$*-lacI (spc)* | SPP1 NRS5742→NRS3936 |
| NRS5770 | 168 *amyE::P-spank-tapA*$_{Bs\_1-188aa}$*-lacI (spc)* | pNW1806→168 |
| NRS5771 | 168 *amyE::P-spank-tapA*$_{Bs\_1-183aa}$*-lacI (spc)* | pNW1807→168 |
| NRS5772 | 168 *amyE::P-spank-tapA*$_{Bs\_1-178aa}$*-lacI (spc)* | pNW1808→168 |
| NRS5785 | 168 *amyE::P-spank-tapA*$_{Bs\_1-173aa}$*-lacI (spc)* | pNW1810→168 |
| NRS5789 | NCIB3610 *ΔtapA + amyE::P-spank-tapA*$_{B\_sub\_1-188aa}$*-lacI (spc)* | SPP1 NRS5770→NRS3936 |
| NRS5790 | NCIB3610 *ΔtapA + amyE::P-spank-tapA*$_{B\_sub\_1-183aa}$*-lacI (spc)* | SPP NRS15771→NRS3936 |
| NRS5791 | NCIB3610 *ΔtapA + amyE::P-spank-tapA*$_{Bs\_1-178aa}$*-lacI (spc)* | SPP1 NRS5772→NRS3936 |
| NRS5793 | NCIB3610 *ΔtapA + amyE::P-spank-tapA*$_{Bs\_1-173aa}$*-lacI (spc)* | SPP1 NRS5785→NRS3936 |
| NRS5799 | 168 *amyE::P-spank-tapA*$_{Bs\_1-133aa}$*-lacI (spc)* | pNW1815→168 |
| NRS5800 | 168 *amyE::P-spank-tapA*$_{Bs\_1-143aa}$*-lacI (spc)* | pNW1816→168 |
| NRS5801 | 168 *amyE::P-spank-tapA*$_{Bs\_1-153aa}$*-lacI (spc)* | pNW1817→168 |
| NRS5802 | 168 *amyE::P-spank-tapA*$_{Bs\_1-163aa}$*-lacI (spc)* | pNW1818→168 |

| Strain | Relevant genotype/description [a] | Source/construction [b] |
|---|---|---|
| NRS5805 | NCIB3610 *ΔtapA + amyE::P-spank-tapA*$_{Bs\_1\text{-}133aa}$*-lacI (spc)* | SPP1 NRS5799→NRS3936 |
| NRS5806 | NCIB3610 *ΔtapA + amyE::P-spank-tapA*$_{Bs\_1\text{-}143aa}$*-lacI (spc)* | SPP1 NRS5800→NRS3936 |
| NRS5813 | NCIB3610 *ΔtapA + amyE::P-spank-tapA*$_{Bs\_1\text{-}153aa}$*-lacI (spc)* | SPP1 NRS5801→NRS3936 |
| NRS5814 | NCIB3610 *ΔtapA + amyE::P-spank-tapA*$_{Bs\_1\text{-}163aa}$*-lacI (spc)* | SPP1 NRS5802→NRS3936 |
| NRS5819 | 168 *amyE::P-spank-tapA*$_{Bs\_1\text{-}123aa}$*-lacI* (spc) | pNW1819→168 |
| NRS5985 | 168 *amyE::P-spank-tapA*$_{Bs\_1\text{-}113aa}$*-lacI (spc)* | pNW1820→168 |
| NRS5986 | 168 *amyE::P-spank-tapA*$_{Bs\_1\text{-}103aa}$*-lacI (spc)* | pNW1821→168 |
| NRS5987 | NCIB3610 *ΔtapA + amyE::P-spank-tapA*$_{Bs\_1\text{-}123aa}$*-lacI (spc)* | SPP1 NRS5819→NRS3936 |
| NRS5988 | NCIB3610 *ΔtapA + amyE::P-spank-tapA*$_{Bs\_1\text{-}113aa}$*-lacI (spc)* | SPP1 NRS5985→NRS3936 |
| NRS5989 | NCIB3610 *ΔtapA + amyE::P-spank-tapA*$_{Bs\_1\text{-}103aa}$*-lacI (spc)* | SPP1 NRS5986→NRS3936 |
| NRS5990 | 168 *bpr::erm trpC2* | BKE15300 (Koo *et al.*, 2017a) |
| NRS5996 | 168 *amyE::P-spank-tapA*$_{Bs\_1\text{-}50aa}$*-lacI (spc)* | pNW1822→168 |
| NRS5997 | 168 *amyE::P-spank-tapA*$_{Bs\_1\text{-}71aa}$*-lacI (spc)* | pNW1823→168 |
| NRS5998 | 168 *amyE::P-spank-tapA*$_{Bs\_1\text{-}88aa}$*-lacI (spc)* | pNW1824→168 |
| NRS5999 | 168 *amyE::P-spank-tapA*$_{Bs\_1\text{-}95aa}$*-lacI (spc)* | pNW1825→168 |
| NRS6002 | NCIB3610 *ΔtapA + amyE::P-spank-tapA*$_{Bs\_1\text{-}50aa}$*-lacI (spc)* | SPP1 NRS5996→NRS3936 |
| NRS6003 | NCIB3610 *ΔtapA + amyE::P-spank-tapA*$_{Bs\_1\text{-}71aa}$*-lacI (spc)* | SPP1 NRS5997→NRS3936 |
| NRS6004 | NCIB3610 *ΔtapA + amyE::P-spank-tapA*$_{Bs\_1\text{-}88aa}$*-lacI (spc)* | SPP1 NRS5998→NRS3936 |
| NRS6005 | NCIB3610 *ΔtapA + amyE::P-spank-tapA*$_{Bs\_1\text{-}95aa}$*-lacI (spc)* | SPP1 NRS5999→NRS3936 |
| NRS6010 | 168 *vpr::erm trpC2* | BKE38090 (Koo et al., 2017a) |
| NRS6011 | 168 *nprB::erm trpC2* | BKE11100 (Koo et al., 2017a) |
| NRS6012 | 168 *mpr::erm trpC2* | BKE02240 (Koo et al., 2017a) |
| NRS6013 | 168 *epr::erm trpC2* | BKE38400 (Koo et al., 2017a) |
| NRS6014 | 168 *aprE::erm trpC2* | BKE10300 (Koo et al., 2017a) |
| NRS6015 | 168 *nprE::erm trpC2* | BKE14700 (Koo et al., 2017a) |
| NRS6016 | *E. coli* CDR 1055 pDR244 | BGSC ECE274 |
| NRS6017 | NCIB3610 *comI*$_{Q12L}$ | BGSC 3A38 |
| NRS6024 | 168 *amyE::P-spank-tapA*$_{Bs\_1\text{-}56aa}$*-lacI (spc)* | pNW1829→168 |
| NRS6025 | NCIB3610 *ΔtapA + amyE::P-spank-tapA*$_{Bs\_1\text{-}56aa}$*-lacI (spc)* | SPP1 NRS6024→NRS3936 |
| NRS6036 | 168 *amyE::P-spank-tapA*$_{Bs\_1\text{-}57aa}$*-lacI (spc)* | pNW1830→168 |
| NRS6037 | 168 *amyE::P-spank-tapA*$_{Bs\_1\text{-}58aa}$*-lacI (spc)* | pNW1831→168 |
| NRS6038 | 168 *amyE::P-spank-tapA*$_{Bs\_1\text{-}59aa}$*-lacI (spc)* | pNW1832→168 |
| NRS6039 | 168 *amyE::P-spank-tapA*$_{Bs\_1\text{-}60aa}$*-lacI (spc)* | pNW1833→168 |
| NRS6040 | 168 *amyE::P-spank-tapA*$_{Bs\_1\text{-}65aa}$*-lacI (spc)* | pNW1834→168 |
| NRS6041 | NCIB3610 *ΔtapA + amyE::P-spank-tapA*$_{Bs\_1\text{-}57aa}$*-lacI (spc)* | SPP1 NRS6036→NRS3936 |
| NRS6042 | NCIB3610 *ΔtapA + amyE::P-spank-tapA*$_{Bs\_1\text{-}58aa}$*-lacI (spc)* | SPP1 NRS6037→NRS3936 |
| NRS6043 | NCIB3610 *ΔtapA + amyE::P-spank-tapA*$_{Bs\_1\text{-}59aa}$*-lacI (spc)* | SPP1 NRS6038→NRS3936 |
| NRS6044 | NCIB3610 *ΔtapA + amyE::P-spank-tapA*$_{Bs\_1\text{-}60aa}$*-lacI (spc)* | SPP1 NRS6039→NRS3936 |
| NRS6045 | NCIB3610 *ΔtapA + amyE::P-spank-tapA*$_{Bs\_1\text{-}65aa}$*-lacI (spc)* | SPP1 NRS6040→NRS3936 |
| NRS6046 | NCIB3610 *comI bpr::erm* | SPP1 NRS6046→NRS6017 |
| NRS6047 | NCIB3610 *comI Δbpr* (KO1) | pDR244→NRS6046 |
| NRS6048 | NICB3610 *comI Δbpr vpr::erm* | gDNA 6010→NRS6047 |
| NRS6049 | NCIB3610 *comI Δbpr Δvpr* (KO2) | pDR244→NRS6048 |
| NRS6060 | NCIB3610 *comI Δbpr Δvpr nprB::erm* | gDNA 6011→NRS6049 |
| NRS6061 | NCIB3610 *comI Δbpr Δvpr ΔnprB* (KO3) | pDR244→NRS6060 |
| NRS6062 | NCIB3610 *comI Δbpr Δvpr ΔnprB mpr::erm* | SPP1 NRS6012→NRS6061 |

| Strain | Relevant genotype/description [a] | Source/construction [b] |
|---|---|---|
| NRS6063 | NCIB3610 *comI Δbpr Δvpr ΔnprB Δmpr* (KO4) | pDR244→NRS6062 |
| NRS6064 | NCIB3610 *comI Δbpr Δvpr ΔnprB Δmpr epr::erm* | gDNA 6013→NRS6063 |
| NRS6065 | NCIB3610 *comI Δbpr Δvpr ΔnprB Δmpr Δepr* (KO5) | pDR244→NRS6065 |
| NRS6340 | NCIB3610 *comI Δbpr Δvpr ΔnprB Δmpr Δepr aprE::erm* | gDNA 6014→NRS6063 |
| NRS6341 | NCIB3610 *comI Δbpr Δvpr ΔnprB Δmpr Δepr ΔaprE* (KO6) | pDR244→NRS6340 |
| NRS6361 | NCIB3610 *comI Δbpr Δvpr ΔnprB Δmpr Δepr ΔaprE nprE::erm* | gDNA 6015→NRS6063 |
| NRS6362 | NCIB3610 *comI Δbpr Δvpr ΔnprB Δmpr Δepr ΔaprE ΔnprE* (KO7) | pDR244→NRS6361 |
| NRS6293 | NCIB3610 *comI Δbpr Δvpr ΔnprB Δmpr Δepr ΔaprE ΔnprE* (KO7) *ΔtapA* | pNW685→NRS6362 |
| NRS6295 | NCIB3610 *comI Δbpr Δvpr ΔnprB Δmpr Δepr ΔaprE ΔnprE* (KO7) *ΔtapA amyE::P-spank-tapA*$_{B\_sub*}$*-lacI (spc)* | SPP1 NRS5041→NRS6293 |
| NRS6533 | NCIB3610 *comI Δbpr Δvpr ΔnprB Δmpr Δepr ΔaprE ΔnprE* wprA::Kan | gDNA BKK10770→NRS6362 (Koo et al., 2017b)) |
| NRS5645 | NCIB3610 *comI Δbpr Δvpr ΔnprB Δmpr Δepr ΔaprE ΔnprE* ΔwprA (KO8) | pDR244→NRS5645 |
| NRS5646 | NCIB3610 *comI* KO8 *ΔtapA* | pNW685→NRS5645 |
| NRS5647 | NCIB3610 *comI* KO8 *ΔtapA amyE::P-spank-tapA*$_{B\_sub*}$*-lacI (spc)* | SPP1 NRS5041→NRS5646 |
| NRS6960 | NCIB3610 *comI* KO8 *amyE::P-spank-tapA*$_{Bs\_1\text{-}57aa}$*-lacI (spc)* | SPP1 NRS6041→NRS5645 |
| NRS6961 | NCIB3610 *comI* KO8 *ΔtapA amyE::P-spank-tapA*$_{Bs\_1\text{-}57aa}$*-lacI (spc)* | SPP1 NRS6041→NRS5646 |
| NRS6366 | 168 *amyE::P-spank-tapA*$_{B\_sub\_1\text{-}57aa(T57A)}$*-lacI (spc)* | pNW1849→168 |
| NRS6367 | 168 *amyE::P-spank-tapA*$_{B\_sub\_1\text{-}57aa(Q56A)}$*-lacI (spc)* | pNW1850→168 |
| NRS6368 | 168 *amyE::P-spank-tapA*$_{B\_sub\_1\text{-}57aa(L55I)}$*-lacI (spc)* | pNW1851→168 |
| NRS6369 | 168 *amyE::P-spank-tapA*$_{B\_sub\_1\text{-}57aa(L55K)}$*-lacI (spc)* | pNW1852→168 |
| NRS6370 | 168 *amyE::P-spank-tapA*$_{B\_sub\_1\text{-}57aa(S54A)}$*-lacI (spc)* | pNW1853→168 |
| NRS6371 | 168 *amyE::P-spank-tapA*$_{B\_sub\_1\text{-}57aa(V53I)}$*-lacI (spc)* | pNW1854→168 |
| NRS6372 | 168 *amyE::P-spank-tapA*$_{B\_sub\_1\text{-}57aa(F51A)}$*-lacI (spc)* | pNW1857→168 |
| NRS6373 | 168 *amyE::P-spank-tapA*$_{B\_sub\_1\text{-}57aa(L55A)}$*-lacI (spc)* | pNW1858→168 |
| NRS6374 | 168 *amyE::P-spank-tapA*$_{B\_sub\_1\text{-}57aa(V53A)}$*-lacI (spc)* | pNW1859→168 |
| NRS6375 | 168 *amyE::P-spank-tapA*$_{B\_sub\_1\text{-}57aa(F45A)}$*-lacI (spc)* | pNW1862→168 |
| NRS6376 | 168 *amyE::P-spank-tapA*$_{B\_sub\_1\text{-}57aa(D47A)}$*-lacI (spc)* | pNW1863→168 |
| NRS6377 | 168 *amyE::P-spank-tapA*$_{B\_sub\_1\text{-}57aa(D52A)}$*-lacI (spc)* | pNW1866→168 |
| NRS6378 | 168 *amyE::P-spank-tapA*$_{B\_sub\_1\text{-}57aa(D52L)}$*-lacI (spc)* | pNW1867→168 |
| NRS6384 | NCIB3610 *ΔtapA + amyE::P-spank-tapA*$_{B\_sub\_1\text{-}57aa(T57A)}$*-lacI (spc)* | SPP1 NRS6366→NRS3936 |
| NRS6385 | NCIB3610 *ΔtapA + amyE::P-spank-tapA*$_{B\_sub\_1\text{-}57aa(Q56A)}$*-lacI (spc)* | SPP1 NRS6367→NRS3936 |
| NRS6386 | NCIB 3610 *ΔtapA + amyE::P-spank-tapA*$_{B\_sub\_1\text{-}57aa(L55I)}$*-lacI (spc)* | SPP1 NRS6368→NRS3936 |
| NRS6387 | NCIB3610 *ΔtapA + amyE::P-spank-tapA*$_{B\_sub\_1\text{-}57aa(L55K)}$*-lacI (spc)* | SPP1 NRS6369→NRS3936 |
| NRS6388 | NCIB3610 *ΔtapA + amyE::P-spank-tapA*$_{B\_sub\_1\text{-}57aa(S54A)}$*-lacI (spc)* | SPP1 NRS6370→NRS3936 |
| NRS6389 | NCIB3610 *ΔtapA + amyE::P-spank-tapA*$_{B\_sub\_1\text{-}57aa(V53I)}$*-lacI (spc)* | SPP1 NRS6371→NRS3936 |
| NRS6390 | NCIB3610 *ΔtapA + amyE::P-spank-tapA*$_{B\_sub\_1\text{-}57aa(F51A)}$*-lacI (spc)* | SPP1 NRS6372→NRS3936 |
| NRS6472 | NCIB3610 *ΔtapA + amyE::P-spank-tapA*$_{B\_sub\_1\text{-}57aa(L55A)}$*-lacI (spc)* | SPP1 NRS6373→NRS3936 |
| NRS6473 | NCIB3610 *ΔtapA + amyE::P-spank-tapA*$_{B\_sub\_1\text{-}57aa(V53A)}$*-lacI (spc)* | SPP1 NRS6374→NRS3936 |
| NRS6474 | NCIB3610 *ΔtapA + amyE::P-spank-tapA*$_{B\_sub\_1\text{-}57aa(F45A)}$*-lacI (spc)* | SPP1 NRS6375→NRS3936 |
| NRS6475 | NCIB3610 *ΔtapA + amyE::P-spank-tapA*$_{B\_sub\_1\text{-}57aa(D47A)}$*-lacI (spc)* | SPP1 NRS6376→NRS3936 |
| NRS6476 | NCIB3610 *ΔtapA + amyE::P-spank-tapA*$_{B\_sub\_1\text{-}57aa(D52A)}$*-lacI (spc)* | SPP1 NRS6377→NRS3936 |
| NRS6477 | NCIB3610 *ΔtapA + amyE::P-spank-tapA*$_{B\_sub\_1\text{-}57aa(D52L)}$*-lacI (spc)* | SPP1 NRS6378→NRS3936 |
| NRS6479 | 168 *amyE::P-spank-tapA*$_{B\_sub\_1\text{-}57aa(V53K)}$*-lacI (spc)* | pNW1865→168 |
| NRS6501 | 168 *amyE::P-spank-tapA*$_{B\_sub\_1\text{-}57aa(D52N)}$*-lacI (spc)* | pNW1869→168 |
| NRS6502 | NCIB3610 *ΔtapA + amyE::P-spank-tapA*$_{B\_sub\_1\text{-}57aa(V53K)}$*-lacI (spc)* | SPP1 NRS6479→NRS3936 |
| NRS6516 | NCIB3610 *ΔtapA + amyE::P-spank-tapA*$_{B\_sub\_1\text{-}57aa(D52N)}$*-lacI (spc)* | SPP1 NRS6501→NRS3936 |

| Strain | Relevant genotype/description [a] | Source/construction [b] |
| --- | --- | --- |
| NRS6520 | NCIB3610 *amyE::P-spank- tapA*$_{B\_sub\_1\text{-}57aa(L55K)}$*-lacI (spc)* | SPP1 NRS6369→NCIB3610 |
| NRS6521 | NCIB3610 *amyE::P-spank- tapA*$_{B\_sub\_1\text{-}57aa(L55A)}$*-lacI (spc)* | SPP1 NRS6373→NCIB3610 |
| NRS6522 | NCIB3610 *amyE::P-spank- tapA*$_{B\_sub\_1\text{-}57aa(F45A)}$*-lacI (spc)* | SPP1 NRS6375→NCIB3610 |
| NRS6535 | 168 *amyE::P-spank- tapA*$_{RBS}$*-tasA*$_{1\text{-}27}$*-tapA*$_{44\text{-}253}$*-lacI (spc)* | pNW1885→168 |
| NRS6536 | NCIB3610 *ΔtapA + amyE::P-spank- tapA*$_{RBS}$*-tasA*$_{1\text{-}27}$*-tapA*$_{44\text{-}253}$*-lacI (spc)* | SPP1 NRS6535→NRS3936 |
| BKK10770 | 168 *wprA::kan trpC2* | BGSC (Koo et al., 2017a) |
| NRS6810 | NCIB3610 *comI ΔwprA* | gDNA BKK10770→NRS6017 |

[a.] Drug resistance cassettes are indicated as the following: (*spc)* spectinomycin resistance, (*erm*) erythromycin resistance and (*cml*) chloramphenicol resistance. *Allele of *tapA* amplified from NCIB3610 (B_sub or Bs), **Allele of *tapA* amplified from *Bacillus pumilis* strain SAFR-032 (B_pum), ***Allele of *tapA* amplified from *Bacillus amyloliquefaciens* strain FZB42 (B_amy) and ****Allele of *tapA* amplified from *Bacillus paralicheniformis* (B_para).

[b.] BGSC is the *Bacillus* genetic stock centre. The direction of strain construction is indicated with an arrow (→) from the genetic source either phage (SPP1), plasmid (pNW) or genomic DNA (gDNA) to the recipient strain.

Table S2 Plasmids used in this study

| Plasmid | Description [a] | Source |
|---|---|---|
| pDR110 | *B. subtilis* integration vector for IPTG-induced expression | (Britton *et al.*, 2002) |
| pGEX-6P-1 | Vector for overexpression of GST-fused proteins | GE Healthcare |
| pMiniMAD | Temp sensitive allelic replacement vector | (Patrick & Kearns, 2008b) |
| pNW685 | pMini-Mad Δ*tapA* | This work (Laura Hobley) |
| pNW1080 | *E. coli* BL21 pGEX-6P-1+TEV-site+ser-$tasA_{B\_sub28\text{-}261aa}$ | (Erskine et al., 2018b) |
| pNW1437 | *E. coli* BL21 pGEX-6P-1+TEV-site+$tasA_{B\_sub28\text{-}261aa}$ | (Erskine et al., 2018b) |
| pNW1448 | pMini-Mad Δ*tasA* | (Erskine et al., 2018b) |
| pNW1438 | pDR110-$tapA_{B\_sub*}$ | This work (Rachel Gillespie) |
| pNW1439 | pDR110-$tapA_{B\_pum**}$ | This work (Rachel Gillespie) |
| pNW1440 | pDR110-$tapA_{B\_amy***}$ | This work (Rachel Gillespie) |
| pNW1800 | pDR110-$tapA_{B\_para****}$ | This work |
| pNW1801 | pDR110-$tapA_{B\_sub\_1\text{-}193aa}$ | This work |
| pNW1806 | pDR110-$tapA_{B\_sub\_1\text{-}188aa}$ | This work |
| pNW1807 | pDR110-$tapA_{B\_sub\_1\text{-}183aa}$ | This work |
| pNW1808 | pDR110-$tapA_{B\_sub\_1\text{-}178aa}$ | This work |
| pNW1810 | pDR110-$tapA_{B\_sub\_1\text{-}173aa}$ | This work |
| pNW1818 | pDR110-$tapA_{B\_sub\_1\text{-}163aa}$ | This work |
| pNW1817 | pDR110-$tapA_{B\_sub\_1\text{-}153aa}$ | This work |
| pNW1816 | pDR110-$tapA_{B\_sub\_1\text{-}143aa}$ | This work |
| pNW1815 | pDR110-$tapA_{B\_sub\_1\text{-}133aa}$ | This work |
| pNW1819 | pDR110-$tapA_{B\_sub\_1\text{-}123aa}$ | This work |
| pNW1820 | pDR110-$tapA_{B\_sub\_1\text{-}113aa}$ | This work |
| pNW1821 | pDR110-$tapA_{B\_sub\_1\text{-}103aa}$ | This work |
| pNW1825 | pDR110-$tapA_{B\_sub\_1\text{-}95aa}$ | This work |
| pNW1824 | pDR110-$tapA_{B\_sub\_1\text{-}88aa}$ | This work |
| pNW1823 | pDR110-$tapA_{B\_sub\_1\text{-}71aa}$ | This work |
| pNW1834 | pDR110-$tapA_{B\_sub\_1\text{-}65aa}$ | This work |
| pNW1833 | pDR110-$tapA_{B\_sub\_1\text{-}60aa}$ | This work |
| pNW1832 | pDR110-$tapA_{B\_sub\_1\text{-}59aa}$ | This work |
| pNW1831 | pDR110-$tapA_{B\_sub\_1\text{-}58aa}$ | This work |
| pNW1830 | pDR110-$tapA_{B\_sub\_1\text{-}57aa}$ | This work |
| pNW1829 | pDR110-$tapA_{B\_sub\_1\text{-}56aa}$ | This work |
| pNW1822 | pDR110-$tapA_{B\_sub\_1\text{-}50aa}$ | This work |
| pNW1849 | pDR110-$tapA_{B\_sub\_1\text{-}57(T57A)}$ | This work |
| pNW1850 | pDR110-$tapA_{B\_sub\_1\text{-}57(Q56A)}$ | This work |
| pNW1851 | pDR110-$tapA_{B\_sub\_1\text{-}57(L55I)}$ | This work |
| pNW1852 | pDR110-$tapA_{B\_sub\_1\text{-}57(L55K)}$ | This work |
| pNW1858 | pDR110-$tapA_{B\_sub\_1\text{-}57(L55A)}$ | This work |
| pNW1853 | pDR110-$tapA_{B\_sub\_1\text{-}57(S54A)}$ | This work |
| pNW1854 | pDR110-$tapA_{B\_sub\_1\text{-}57(V53I)}$ | This work |
| pNW1865 | pDR110-$tapA_{B\_sub\_1\text{-}57(V53K)}$ | This work |
| pNW1859 | pDR110-$tapA_{B\_sub\_1\text{-}57(V53A)}$ | This work |
| pNW1866 | pDR110-$tapA_{B\_sub\_1\text{-}57(D52A)}$ | This work |
| pNW1867 | pDR110-$tapA_{B\_sub\_1\text{-}57(D52L)}$ | This work |
| pNW1869 | pDR110-$tapA_{B\_sub\_1\text{-}57(D52N)}$ | This work |
| pNW1857 | pDR110-$tapA_{B\_sub\_1\text{-}57(F51A)}$ | This work |
| pNW1863 | pDR110-$tapA_{B\_sub\_1\text{-}57(D47A)}$ | This work |
| pNW1862 | pDR110-$tapA_{B\_sub\_1\text{-}57(F45A)}$ | This work |
| pNW1885 | pDR110-$tapA_{RBS}$-$tasA_{1\text{-}27}$-$tapA_{44\text{-}253}$ | Synthesised by DCBiosciences |
| pNW1600 | pGEX-6P-1-$tapA_{B\_sub44\text{-}253}$ | This work |

| Plasmid | Description [a] | Source |
|---|---|---|
| pNW1600 | pGEX-6P-1+TEV-site+*tapA*$_{B\_sub44-253aa}$ | This work (Laura D'Ignazio) |
| pNW1441 | pGEX-6P-1+TEV-site+*tapA*$_{B\_sub34-253aa}$ | This work (Rachel Gillespie) |
| pNW2021 | pMini-Mad Δ*sipW* | Synthesised by GenScript (Elliot Erskine) |

[a.] *Allele of *tapA* amplified from NCIB 3610 (B_sub), **Allele of *tapA* amplified from *Bacillus pumilis* strain SAFR-032 (B_pum), ***Allele of *tapA* amplified from *Bacillus amyloliquefaciens* strain FZB42 (B_amy) and ****Allele of *tapA* amplified from *Bacillus paralicheniformis* (B_para).

| Primer | Sequence 5’ – 3’[a] | Use | Used to make plasmid[b] |
| --- | --- | --- | --- |
| NSW2181 | GCTAGGATCCGAAAATTTATATTTTCAAGCTTTTCATG ATATTGAAACATTTG | Forward primer from TapA$_{\text{B_sub44aa}}$ | pNW1600<br>For:2181<br>Rev:1895<br><br>TapA44-253 into pGEX-6P-1 |
| NSW 1895 | GCATCTCGAG**TTATTA**CTGATCAGCTTCATTGC | Reverse primer from TapA$_{\text{B_sub253aa}}$ | |
| NSW1896 | ATGCGTCGACTTTTACAGGAGGTAAGATATGTTTCG | Forward primer TapA$_{\text{B_sub*}}$ | pNW1438<br>For:1896<br>Rev:1897 |
| NSW1897 | ATGCGCATGC**TTA**CTGATCAGCTTCATTGC | Forward primer TapA$_{\text{B_sub*_full-length}}$ | |
| NSW2147 | GCATCTCTTTCACATCAGTGATTTC | Upstream *sipW* | Checking Δ*sipW* strain |
| NSW2148 | GCATGGGGAAGAGGATGAAAAAA | Downstream *sipW* | |
| NSW2511 | ATGCGCATGCTTAGCATTTTGCAAGCCTCATAGGC | Reverse primer TapA$_{\text{B_sub_1-188aa}}$ | pNW1806<br>Primers<br>For:1896<br>Rev:2511 |
| NSW2512 | ATGCGCATGCTTACATAGGCTCCGACCACTCA | Reverse primer TapA$_{\text{B_sub_1-183aa}}$ | pNW1807<br>Primers<br>For:1896<br>Rev:2512 |
| NSW2513 | ATGCGCATGCTTACTCAAATGTACTGCCGTTT | Reverse primer TapA$_{\text{B_sub_1-178aa}}$ | pNW1808<br>Primers<br>For:1896<br>Rev:2513 |
| NSW2514 | ATGCGCATGCTTAGTTTGCCGGGTAGCCTGCCGGT | Reverse primer TapA$_{\text{B_sub_1-173aa}}$ | pNW1810<br>Primers<br>For:1896<br>Rev:2514 |
| NSW2521 | ATGCGCATGCTTAGATCACGTTCCCATCCTTTAACG | Reverse primer TapA$_{\text{B_sub_1-133aa}}$ | pNW1815<br>Primers<br>For:1896<br>Rev:2521 |
| NSW2520 | ATGCGCATGCTTAGCCGATTTGATTGGAGAC | Reverse primer TapA$_{\text{B_sub_1-143aa}}$ | pNW1816<br>Primers<br>For:1896<br>Rev:2520 |
| NSW2519 | ATGCGCATGCTTATTTCTTGGTCTCAATTTTATAAAG | Reverse primer TapA$_{\text{B_sub_1-153aa}}$ | pNW1817<br>Primers<br>For:1896<br>Rev:2519 |
| NSW2518 | ATGCGCATGCTTATTTAAATGCATAAATGCCGG | Reverse primer TapA$_{\text{B_sub_1-163aa}}$ | pNW1818<br>Primers<br>For:1896<br>Rev:2518 |

| | | | |
|---|---|---|---|
| NSW2522 | ATGCGCATGCTTAGGCATTTTCAAGCTTATGAAGC | Reverse primer TapA$_{B_sub_1-123aa}$ | pNW1819<br>Primers<br>For:1896<br>Rev:2522 |
| NSW2523 | ATGCGCATGCTTACCACTTTGATTTCTTAAGTTTCTCACC | Reverse primer TapA$_{B_sub_1-113aa}$ | pNW1820<br>Primers<br>For:1896<br>Rev:2523 |
| NSW2524 | ATGCGCATGCTTAATTTTCGAGCACAGCAAATAAGGCG | Reverse primer TapA$_{B_sub_1-103aa}$ | pNW1821<br>Primers<br>For:1896<br>Rev:2524 |
| NSW2531 | ATGCGCATGCTTATGTTTCAATATCATGAAAAGC | Reverse primer TapA$_{B_sub_1-50aa}$ | pNW1822<br>Primers<br>For:1896<br>Rev:2531 |
| NSW2532 | ATGCGCATGCTTAATCATAATGGCAGTTTTTATC | Reverse primer TapA$_{B_sub_1-71aa}$ | pNW1823<br>Primers<br>For:1896<br>Rev:2532 |
| NSW2533 | ATGCGCATGCTTATTTCGTATCCGTTTGATCTG | Reverse primer TapA$_{B_sub_1-88aa}$ | pNW1824<br>Primers<br>For:1896<br>Rev:2533 |
| NSW2534 | ATGCGCATGCTTAGAAAGGTGAGCATACAGTGC | Reverse primer TapA$_{B_sub_1-95aa}$ | pNW1825<br>Primers<br>For:1896<br>Rev:2534 |
| NSW2550 | ATGCGCATGCTTATTGAAGTGAGACATCAAATGTTTC | Reverse primer TapA$_{B_sub_1-56aa}$ | pNW1829<br>Primers<br>For:1896<br>Rev:2550 |
| NSW2565 | ATGCGCATGCTTACGTTTGAAGTGAGACATC | Reverse primer TapA$_{B_sub_1-57aa}$ | pNW1830<br>Primers<br>For:1896<br>Rev:2565 |
| NSW2566 | ATGCGCATGCTTAACACGTTTGAAGTGAGAC | Reverse primer TapA$_{B_sub_1-58aa}$ | pNW1831<br>Primers<br>For:1896<br>Rev:2566 |
| NSW2567 | ATGCGCATGCTTATTTACACGTTTGAAGTGAG | Reverse primer TapA$_{B_sub_1-59aa}$ | pNW1832<br>Primers<br>For:1896<br>Rev:2567 |
| NSW2568 | ATGCGCATGCTTAGTCTTTACACGTTTGAAGTGAG | Reverse primer TapA$_{B_sub_1-60aa}$ | pNW1833<br>Primers<br>For:1896<br>Rev:2568 |
| NSW2569 | ATGCGCATGCTTAATCTGTATGCTGAAAGTC | Reverse primer TapA$_{B_sub_1-65aa}$ | pNW1834<br>Primers<br>For:1896<br>Rev:2569 |

| | | | |
|---|---|---|---|
| NSW2001 | ATGCGTCGACCAAGGAGTTGGAGAATGAATGAAACAGTCGC | Forward primer TapA$_{B_pum**}$ | pNW1827 Primers For:2001 Rev:2002 |
| NSW2002 | ATGCGCATGC**TTA**AGATACCTTTTCTGGACACTTTGC | Reverse primer TapA$_{B_pum**_Full-length}$ | |
| NSW2506 | ATGCGTCGACCTCGCTAGGAGGGTAGTCATGTTCC | Forward primer TapA$_{B_para****}$ | pNW1800 Primers For:2506 Rev:2507 |
| NSW2507 | ATGCGCATGCTCATGACGCTTCCCCGCTTTC | Reverse primer TapA$_{B_para\ ****full-length}$ | |
| NSW1898 | gcatGTCGACTTTTACAGGGGGTAAGGCATGTTCCGATTGTTGC | Forward primer TapA$_{B_amy***}$ | pNW1440 Primers For:1898 Rev:1899 |
| NSW1899 | atgcGCATGC**TTA**CTGATCAGTTTCAGCGTTTTTTTCATGTTCTTCC | Reverse primer TapA$_{B_amy***\ full-lenght}$ | |
| NSW2841 | ATGCGCATGC**TTA**AGCTTGAAGTGAGACATC | TapA$_{B_sub_1-57(T57A)}$ | pNW1849 For:1896 Rev:2841 |
| NSW2842 | ATGCGCATGC**TTA**CGTTGCAAGTGAGACATC | TapA$_{B_sub_1-57\ (Q56A)}$ | pNW1850 For:1896 Rev:2842 |
| NSW2843 | ATGCGCATGC**TTA**CGTTTGAATTGAGACATC | TapA$_{B_sub_1-57(L55I)}$ | pNW1851 For:1896 Rev:2843 |
| NSW2844 | ATGCGCATGC**TTA**CGTTTGCTTTGAGACATC | TapA$_{B_sub_1-57(L55K)}$ | pNW1852 For:1896 Rev:2844 |
| NSW2845 | ATGCGCATGC**TTA**CGTTTGAAGTGCGACATC | TapA$_{B_sub_1-57(S54A)}$ | pNW1853 For:1896 Rev:2845 |
| NSW2846 | ATGCGCATGC**TTA**CGTTTGAAGTGAGATATC | TapA$_{B_sub_1-57(V53I)}$ | pNW1854 For:1896 Rev:2846 |
| NSW2858 | GATATTGAAACATTTGATAAGTCACTTCAAACGTG | TapA$_{B_sub_1-253(V53K)}$ | pNW1855 For:2858 Rev: 2859 KOD (QC) |
| NSW2859 | CACGTTTGAAGTGACTTATCAAATGTTTCAATATC | TapA$_{B_sub_1-253(V53K)}$ | |
| NSW2849 | ATGCGCATGC**TTA**CGTTTGTGCTGAGACATC | TapA$_{B_sub_1-57(L55A)}$ | pNW1858 For:1896 Rev:2846 |
| NSW2565 | ATGCGCATGC**TTA**CGTTTGAAGTGAGACATC | TapA$_{B_sub_1-57(F51A)}$ | pNW1857 For:1896 Rev:2565 |
| NSW2850 | ATGCGCATGC**TTA**CGTTTGAAGTGATGCATC | TapA$_{B_sub_1-57(V53A)}$ | pNW1859 For:1896 Rev:2850 |
| NSW2854 | GATATTGAAACATTTGCTGTCTCACTTCAAACG | TapA$_{B_sub_1-253(D52A)}$ | pNW1856 For:2854 Rev:2855 KOD (QC) |
| NSW2855 | CGTTTGAAGTGAGACAGCAAATGTTTCAATATC | TapA$_{B_sub_1-253(D52A)}$ | |

| | | | |
|---|---|---|---|
| NSW2584 | GCGCTGCTTTTCATGCTATTGAAACATTTGATGTC | TapA$_{B_sub_1\text{-}253(D47A)}$ | pNW1861 For:2584 Rev:2585 KOD (QC) |
| NSW2585 | GACATCAAATGTTTCAATAGCATGAAAAGCAGCGC | TapA$_{B_sub_1\text{-}253(D47A)}$ | |
| NSW2856 | GATATTGAAACATTTAATGTCTCACTTCAAACG | TapA$_{B_sub_1\text{-}253(D52N)}$ | pNW1868 For:2584 Rev:2585 KOD (QC) |
| NSW2857 | CGTTTGAAGTGAGACTAAAAATGTTTCAATATC | TapA$_{B_sub_1\text{-}253(D52N)}$ | |
| NSW2829 | GATACAAGCGCTGCTGCTCATGATATTGAAACA | TapA$_{B_sub_1\text{-}253(F45A)}$ | pNW1845 For:2829 Rev:2830 KOD (QC) |
| NSW2830 | TGTTTCAATATCATGAGCAGCAGCGCTTGTATC | TapA$_{B_sub_1\text{-}253(F45A)}$ | |
| NSW872 | AGGTGTGGCATAATGTGTGTAATTGTGAGC | pDR110 Forward | Sequencing primers for pDR110 |
| NSW873 | TGAACAATCACGAAACAATAATTGGTACGTACG | pDR110 Reverse | |
| NSW2750 | ACTCCTACGGGAGGCAGC | 16S rRNA gene forward | 16S amplification for phylogeny assignment |
| NSW2751 | TCACGACACGAGCTGACGAC | 16S rRNA gene reverse | |
| NSW1308 | GCATGGATCCCTCTTCCCATTTGGACATGTG | *tapA* upstream for | Primers for the *tapA* deletion construct in the pMAD vector pNW685; also used for screening for mutations on the chromosome. |
| NSW1332 | GGTAAGATATGTTTCGATTGGTCGACATGC | *tapA* upstream rev | |
| NSW1333 | GCATGTCGACCAGAAGGAAAGCGGGGAAGAG | *tapA* downstream for | |
| NSW1334 | GCATGAATTCATATCGAAACCTGTTGCCAGG | *tapA* downstream rev | |

[a.] Restriction sites are underlined, stop codons are in bold.

[b.] QC = QuickChange (using KOD polymerase) and SDM = Site-Directed Mutagenesis.

*Allele of *tapA* amplified from NCIB 3610 (B_sub)

**Allele of *tapA* amplified from *Bacillus pumilis* strain SAFR-032 (B_pum)

***Allele of *tapA* amplified from *Bacillus amyloliquefaciens* strain FZB42 (B_amy)

****Allele of *tapA* amplified from *Bacillus paralicheniformis*.

A

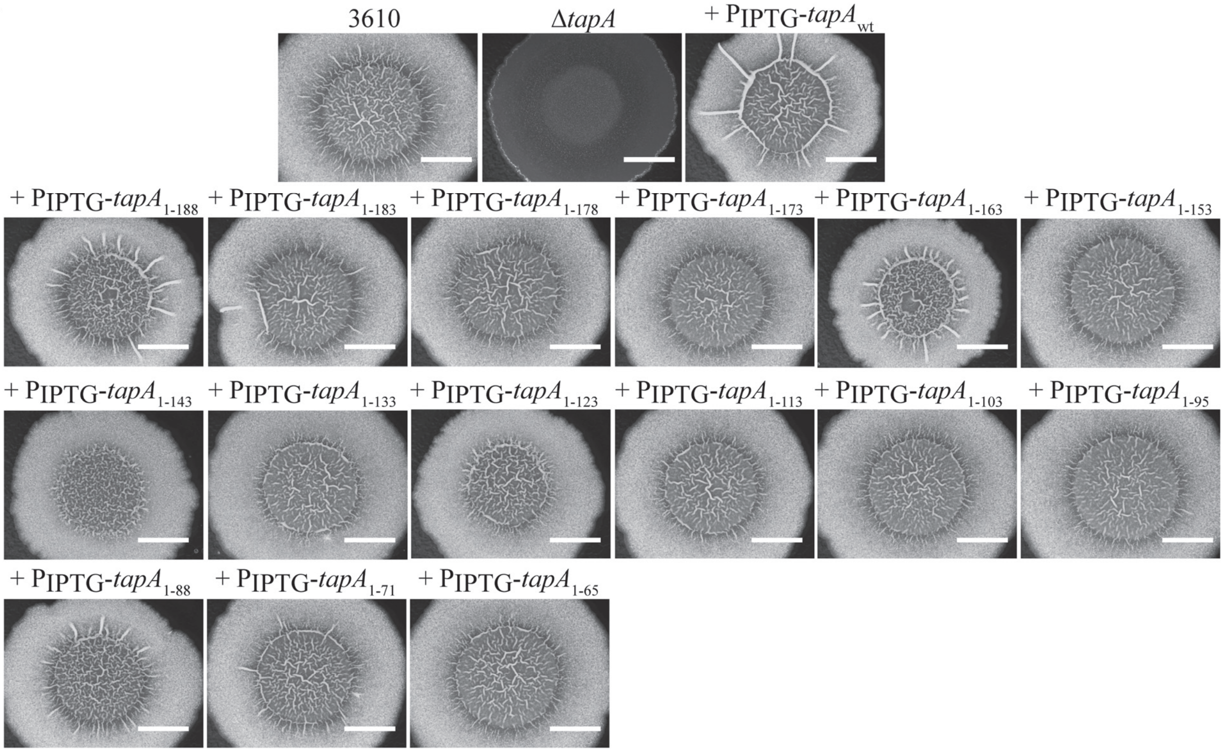


B

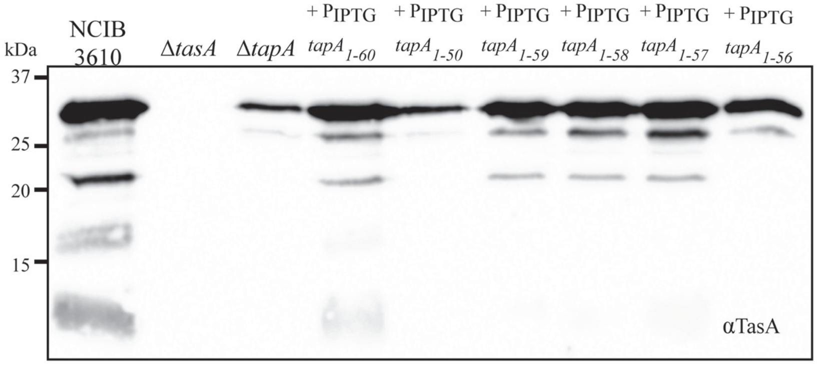


**Figure S1: The *tapA* coding region is functional when truncated.** (**A**) Biofilms formed by NCIB3610, Δ*tapA* (NRS3936), +$P_{IPTG}$-*tapA* (NRS5045), +$P_{IPTG}$-*tapA*$_{1-188}$ (NRS5789), +$P_{IPTG}$-*tapA*$_{1-183}$ (NRS5790), +$P_{IPTG}$-*tapA*$_{1-178}$ (NRS5791), +$P_{IPTG}$-*tapA*$_{1-173}$ (NRS5793), +$P_{IPTG}$-*tapA*$_{1-163}$ (NRS5814), +$P_{IPTG}$-*tapA*$_{1-153}$ (NRS5813), +$P_{IPTG}$-*tapA*$_{1-143}$ (NRS5806), +$P_{IPTG}$-*tapA*$_{1-133}$ (NRS5805), +$P_{IPTG}$-*tapA*$_{1-123}$ (NRS5987), +$P_{IPTG}$-*tapA*$_{1-113}$ (NRS5988), +$P_{IPTG}$-*tapA*$_{1-103}$ (NRS5989), +$P_{IPTG}$-*tapA*$_{1-95}$ (NRS6005), +$P_{IPTG}$-*tapA*$_{1-88}$ (NRS6004), +$P_{IPTG}$-*tapA*$_{1-71}$ (NRS6003), and +$P_{IPTG}$-*tapA*$_{1-65}$ (NRS6045). Biofilms were grown at 30$^{o}$C for 48 hours in the presence of 25 μM IPTG. Biofilm images are representative of at least 3 independent replicates. The scale bars represents 1 cm; (**B**) Immunoblot analysis of proteins extracted from biofilms formed by NCIB3610, Δ*tasA* (NRS5267); Δ*tapA* (NRS3936), $P_{IPTG}$-*tapA*$_{1-60}$ (NRS6044), *tapA*$_{1-50}$ (NRS6002), +$P_{IPTG}$-*tapA*$_{1-59}$

(NRS6043), +$P_{IPTG}$-*tapA*$_{1\text{-}58}$ (NRS6042), +$P_{IPTG}$-*tapA*$_{1\text{-}57}$ (NRS6041), and +$P_{IPTG}$-*tapA*$_{1\text{-}56}$ (NRS6025) using αTasA antibodies. n = 2.

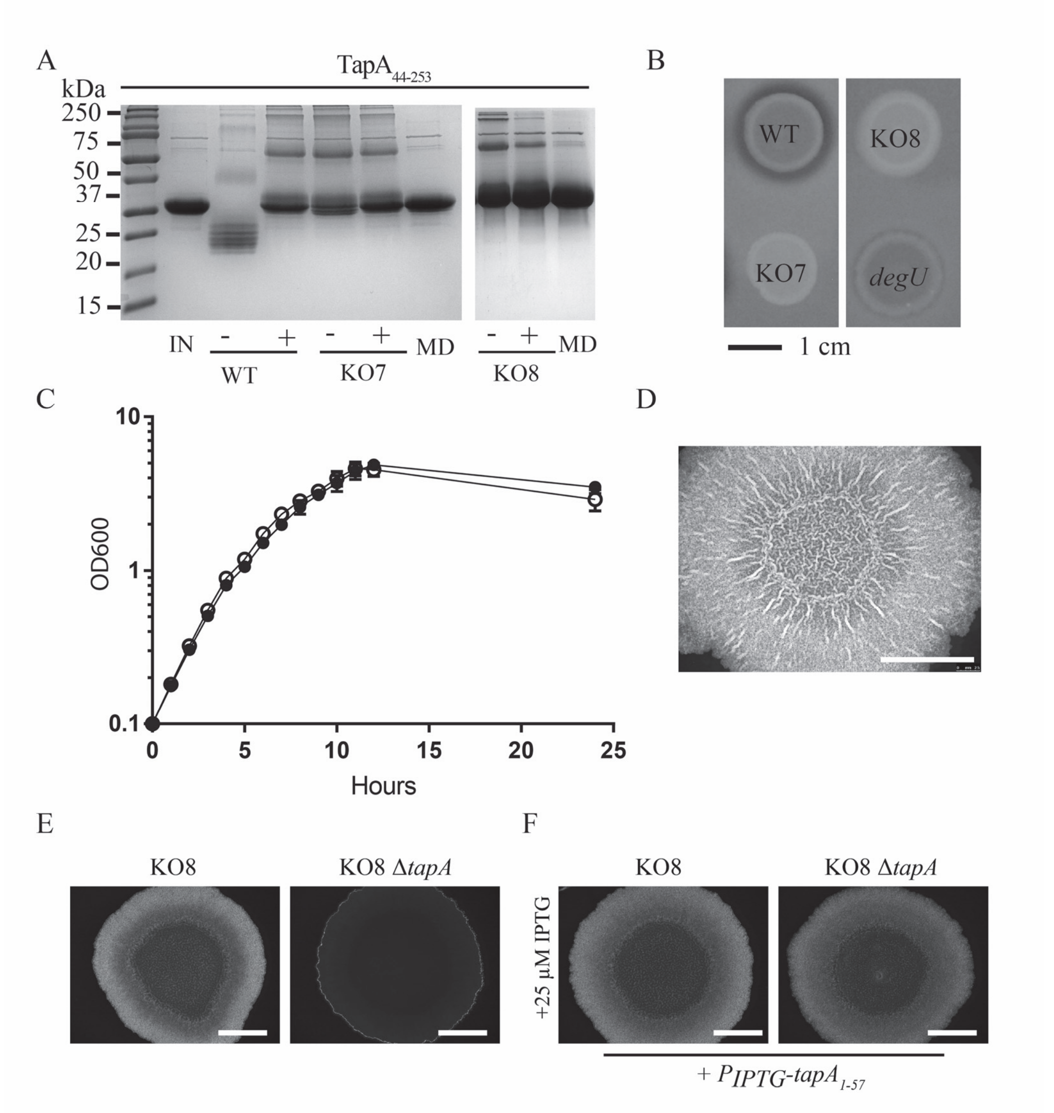


**Figure S2: Recombinant TapA is sensitive to the extracellular proteases secreted by *B. subtilis*.** (**A**) Integrity of 28 µg recombinant TapA$_{44\text{-}253}$ incubated for 8 hrs at 37°C analysed by SDS-PAGE. The protein (IN) was incubated with filtered spent supernatants, or MSgg medium (MD), collected from NCIB3610 *comI* (NRS6017), KO7 (NRS6362) and KO8 (NRS5645). The (-) and (+) indicate if the supernatant had been heat inactivated at 100°C prior to incubation with the recombinant protein. n= 2; (**B**) The ability of strains NCIB3610 *comI*, KO7 (NRS6362), KO8 (NRS5645) and *degU* (NRS1314) to digest milk contained in an agar plates was tested and the results imaged after 20 hours incubation at 37$^{o}$C. n= 2; (**C**) Growth of NCIB3610 *comI* and NCIB3610 *comI* KO8 (NRS5645) in MSgg medium with shaking at 30$^{o}$C measured by OD$_{600}$. An average of two independent experiments are shown as an average with the error bars being the standard deviation; (**D**) Biofilms formed by *B. subtilis* isolates Δ*wprA* (NRS6810); **(E)** NCIB3610 *comI*

KO8 (NRS5645), NCIB3610 *comI* KO8 Δ*tapA* (NRS5646); (**F**) NCIB3610 *comI* KO8 +$P_{IPTG}$-*tapA*$_{1\text{-}57}$ and NCIB3610 *comI* KO8 Δ*tapA* +$P_{IPTG}$-*tapA*$_{1\text{-}57}$ formed in the presence or absence of 25 μM IPTG after growth at 30$^{o}$C for 48 hours. Images representative of 2 independent replicates. The scale bar represents 1 cm.

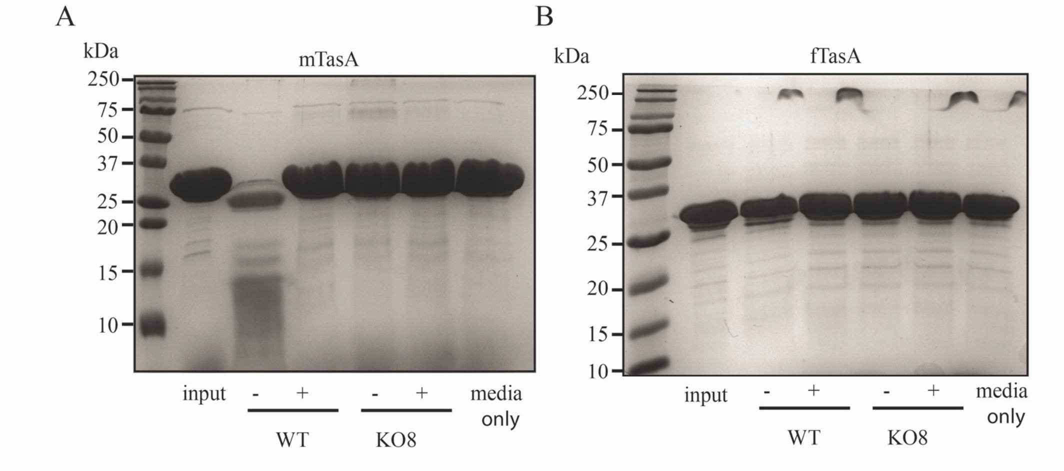


**Figure S3: Recombinant monomeric TasA is sensitive to the extracellular proteases secreted by *B. subtilis*.** Integrity of 28 µg recombinant (**A**) monomeric TasA or (**B**) fibrous TasA incubated for 8 hrs at 37°C analysed by SDS-PAGE. The protein (input) was incubated with filtered spent supernatants collected from NCIB3610, KO7 (NRS6362) and KO8 (NRS5645). The (-) and (+) indicate if the supernatant had been heat inactivated at 100°C prior to incubation with the recombinant protein. A media only control is shown. n= 2.
